# Induction of intracellular wild-type p53 amyloids leading to cellular transformation and tumor formation in mice

**DOI:** 10.1101/2020.06.04.133124

**Authors:** Ambuja Navalkar, Satyaprakash Pandey, Namrata Singh, Amit Kumar Dey, Sandhini Saha, Komal Patel, Bhabani Mohanty, Sachin Jadhav, Pradip Chaudhari, Tushar K. Maiti, Samir K. Maji

**Author notes:** Correspondence: Prof. Samir K. Maji, Department of Biosciences and Bioengineering, IIT Bombay, Powai, Mumbai 400 076, India.

## Abstract

Tumor suppressor p53 mutations, with subsequent loss-of-tumor suppressive function and gain-of oncogenic functions, are associated with more than 50% of human cancers. Aggregation and amyloid formation are also mechanisms by which wild type and mutant p53 might be involved in cancer, but the direct evidence of how aggregated p53 acts as an oncogene is lacking. In this study, we directly demonstrate that wild-type p53 amyloid formation imparts oncogenic properties to normal cells. Cells with p53 amyloids show enhanced survival, apoptotic resistance with increased proliferation and migration rates. The tumorigenic potential of p53 amyloid transformed cells is further confirmed in a mice xenograft model, wherein the tumor showed p53 amyloid aggregates. Gene-expression analysis and proteomic profiling suggest that p53 amyloid formation triggers aberrant expression of pro-oncogenes while downregulating the tumor-suppressive genes. Interestingly, disaggregating p53 rescues the cellular transformation and also inhibits tumor development in mice. We propose that wild-type p53 amyloid formation can potentially contribute to the initiation of tumor development.

## Introduction

p53 has been extensively studied in the field of cancer biology due to its important role in the regulation of anti-proliferative and tumor suppressive programs by activating or repressing key effector genes (Aubrey et al., 2018; Lane and Crawford, 1979; Levine, 1997; Mello and Attardi, 2018; Vousden, 2002). Many studies have convincingly established the mechanism of loss of tumor suppressive functions by p53 and the consequent initiation of cancer (Kim et al., 2009; Moll and Schramm, 1998). The loss of p53 function is mostly associated with its mutations, due to which p53 either misfolds (structural mutation) or is unable to bind to its cognate DNA sequence (contact mutation) (Joerger et al., 2006; Olivier et al., 2010). Mutant p53 proteins not only lose their native tumor suppressive functions but often gain additional oncogenic functions and essentially behave as an oncogene (Freed-Pastor and Prives, 2012; Oren and Rotter, 2010; Rivlin et al., 2011). p53 mutations enhance the proliferative and tumorigenic potential of the cells contributing to various stages of tumor progression and increased anti-cancer drug resistance (Bargonetti and Prives, 2019; Stein et al., 2019). Hotspot mutations in p53 show a common tendency to destabilize/misfold the native conformation of the protein (Wilcken et al., 2012), leading to the formation of aggregates of inactive p53, which are sequestered and targeted by the proteasomal degradation machinery (Ano Bom et al., 2012; Levy et al., 2011). Interestingly, wild-type p53 has also been reported to aggregate *in vitro* (Bullock et al., 1997; Wang and Fersht, 2012, 2015) and also in the cytoplasm and/or nucleus of cancer tissues (De Smet et al., 2017; Moll et al., 1996; Ostermeyer et al., 1996). Recently, many studies have suggested that p53 aggregation and amyloid formation might be the causative factors of p53 loss/gain-of-function in cancers (Ghosh et al., 2017; Lasagna-Reeves et al., 2013; Rangel et al., 2014; Silva et al., 2014). This is further supported by the detection of p53 amyloids in cancer cells and tumor tissue biopsies (Ano Bom et al., 2012; Ghosh et al., 2017; Levy et al., 2011). In addition, we have recently demonstrated that p53 amyloid formation leads to loss of p53 functions such as apoptosis, cell cycle arrest in response to DNA damage, and also induces gain of tumorigenic functions. Moreover, similar to many other amyloids associated with neurodegenerative disorders, p53 amyloids also showed prion-like properties in cells (Ghosh et al., 2017). In this study, we ask whether the wild-type p53 amyloid formation can induce the cancerous transformation of normal cells leading to tumor initiation.

Here, we show that the exogenous addition of *in vitro* synthesized p53 core fibrils (seeds) to normal cells induces aggregation of the endogenous p53 in cells. These cells with p53 amyloids exhibit enhanced cancer-reminiscent properties like increased survival, colony formation and proliferation rate, enhanced migration with resistance to apoptosis. More importantly, p53 amyloid containing cells are tumorigenic and can establish a tumor xenograft in immunocompromised mice. Gene expression analysis along with proteomic profiling of the p53 amyloid containing cells indicates the downregulation of anti-cancer genes with a concomitant upregulation of pro-cancer genes involved in the cell cycle, cellular proliferation and signaling, promoting the cancer-like phenotype. Dis-aggregation of cellular p53 aggregates, using a peptide inhibitor rescues p53 functions and reduced the tumorigenic properties of these cells. Overall, our study reveals that wild-type p53 amyloid formation can initiate cancer-like properties contributing to the disease pathology via a collateral loss of tumor suppressive function and a gain-of-oncogenic-function mechanism. These results establish p53 amyloid formation as a plausible cause of cancer initiation.

## Results

### p53 amyloid formation leads to loss of tumor suppressive apoptotic function along with increased oncogenic potential

To examine whether wild-type p53 amyloid formation can lead to the transformation of normal cells to a cancerous phenotype, we chose MCF 10A cells (non-tumorigenic human breast epithelial cells) and HFF cells (normal human foreskin fibroblast). These cells have been used to identify oncogenic agents, which can cause transformation in cells (Imbalzano et al., 2009; Scott et al., 2004; Yusuf and Frenkel, 2010). Since exogenous addition of p53 core amyloid fibrils (seeds) leads to endogenous p53 amyloid formation in cells (Ghosh et al., 2017), we have used this method to induce p53 amyloid formation in these two normal cells. For amyloid seed preparation, p53 core protein was incubated and its aggregation and amyloid formation was confirmed using Circular Dichroism (CD), Thioflavin T (ThT) fluorescence and electron microscopic imaging (Figure 1A, S1A-C). Subsequently, MCF 10A and HFF cells were treated with sonicated core p53 fibrils for 48 hrs to induce aggregation of p53 and untreated cells were kept as control (Figure S1D). Immunofluorescence using p53 DO-1 antibody and confocal microscopy confirm p53 punctate formation in the cells (Figure 1B, S1E). Interestingly, the majority of the cell population showed the presence of nuclear aggregates of p53, both for MCF 10A (~79%) and HFF cells (~83%) (Figure S1F). p53 core monomer treated cells (data not shown), untreated cells and α-synuclein amyloid fibril treated cells did not show significant p53 stabilization (Figure S1E), indicating that p53 aggregation in cells is specific to the treatment of p53 core derived amyloids. To confirm the amyloid nature of p53 aggregates in cells, p53 was immunoprecipitated using p53 DO-1 antibody from core fibril treated and untreated cells (MCF 10A and HFF), followed by dot blot with p53 DO-1 and amyloid fibril specific OC antibody. The dot blot confirms the amyloid nature of p53 aggregates in the fibril treated MCF 10A and HFF cells whereas a significant level of p53 was not detected in untreated cells (Figure 1C, S1G).

**Figure 1.**
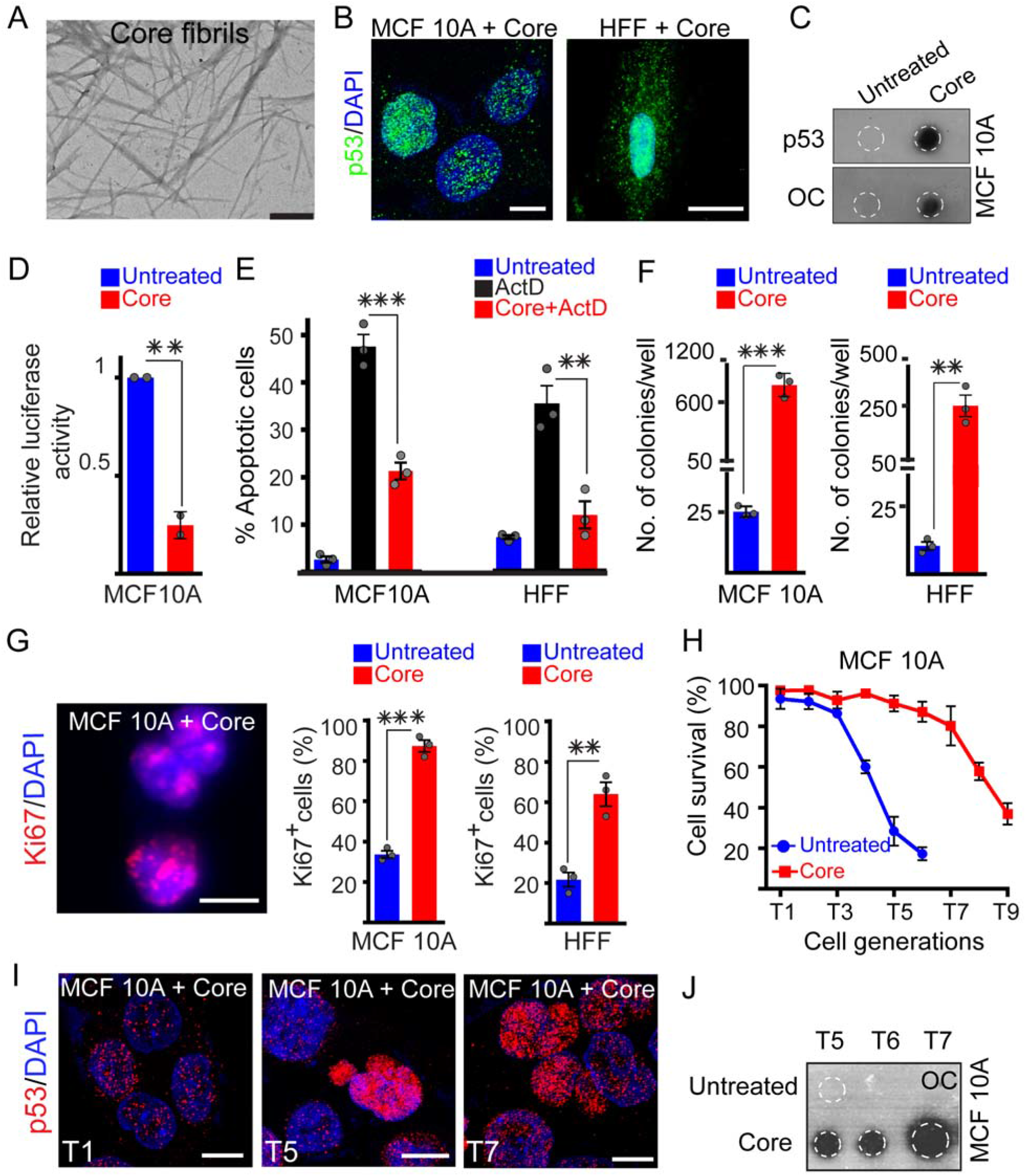
p53 amyloid formation in cells and its consequences. (A) Electron micrograph (EM) of p53 core domain aggregates formed in presence of chondroitin sulphate A (CSA) showing the morphology of amyloid fibrils. Scale bars are 500 nm. n=3 independent experiments. (B) Immunostaining using p53 antibody (DO-1) of MCF 10A and HFF cells that are treated with p53 fibril seeds showing the presence of p53 punctate, predominantly in the nucleus. Scale bar is 10 μm. n=3 independent experiments. (C) Immunoprecipitation of p53 from the MCF 10A core fibril treated cells and subsequent dot blot analysis using amyloid fibril specific antibody (OC) showing the presence of p53 amyloid. The untreated cells did not show the presence of p53 amyloid. n=2 independent experiments. (D) Luciferase reporter assay examining the transcriptional activity of p53 in MCF 10A core fibril treated cells. The core fibril treated cells showing a decrease in luciferase activity compared to control cells (without fibril treatment) indicating loss of p53 binding to its response element. The values plotted represent mean ± s.d, n=2 independent experiments. The p-value of core treated cells with respect to untreated cells is 0.00417. (E) Loss of apoptotic function under stress conditions due to p53 amyloid formation. Core fibril treated MCF 10A and HFF cells showing significantly less apoptosis under ActD treatment (early and late combined) compared to cells that received only ActD treatment. For (E), values represent mean ± s.e.m, n=3 independent experiments. The p-value for MCF 10A is 0.0010 and HFF is 0.0075, calculated with respect to only ActD treated cells. (F) Effect of p53 amyloid formation on transforming ability of MCF 10A and HFF cells. Soft agar colony formation assay of both the cells showing significantly higher colony formation when pre-treated with core fibril seeds as compared to the untreated control. The data suggesting upon p53 amyloid formation, both the cells gain enhanced transforming properties. The values were plotted representing mean ± s.e.m, n=3 independent experiments. The p values for MCF 10A and HFF core treated cells with respect to untreated cells are 0.00028 and 0.00323, respectively. (G) Effect of p53 amyloid formation on the proliferation of cells. Immunostaining of core fibril treated MCF 10A cells (left panel) showing expression of Ki67 (proliferation marker) with nuclear localization of Ki67. Quantification of Ki67 positive (Ki67^+^) cells showing a higher Ki67^+^ population for fibril treated MCF 10A (middle panel) and HFF cell (right panel) compared to untreated control. Scale bars are 10 μm. For (G), values represent mean ± s.e.m, n=3 independent experiments. The p-value for MCF10A core treated cells is <0.0001 and for HFF core treated cells is 0.00346, calculated against respective untreated controls. (H) Inter-generation transmission of p53 amyloid aggregates. MCF 10A were treated with core fibrils and continuously passaged for evaluating the survival as compared to untreated cells (also correspondingly passaged). Each cell generation was labeled as T1, T2 and so on. Cell viability of fibril treated and untreated cells was calculated using Annexin-V PI assay coupled with FACS analysis. Untreated cells showing decreased viability at higher passage number as compared to the fibril treated cells. The values in the plots represent mean ± s.d, n=2 independent experiments. (I) Immunofluorescence staining of core fibril treated MCF 10A cells using p53 DO-1 antibody showing the presence of p53 aggregates in the T1, T5 and T7 passages. Scale bars are 10 μm. (J) Immunoprecipitation of p53 aggregates from MCF 10A cells followed by dot blot analysis using OC (amyloid fibril specific) antibodies showing the presence of p53 fibrils in the core treated cells for various generations. Untreated cells did not show the presence of p53 amyloid aggregates. Statistical significance (***p ≤ 0.001, **p ≤ 0.01, *p ≤ 0.05) is determined by one-way ANOVA followed by Bonferroni multiple comparison post hoc test with 95% confidence interval.

For tumor suppressive functions, p53 modulates cellular expression of several hundreds of genes by binding to its specific DNA Response Elements (Kastenhuber and Lowe, 2017). Since p53 formed aggregates inside the nucleus of cells, we first examined this specific DNA binding and downstream transcriptional activity of p53 amyloids in MCF 10A cells using a luciferase reporter assay (El-Deiry et al., 1993). Core fibril treated cells showed significant loss of p53 transcriptional activity (luciferase signal) as compared to the untreated cells, suggesting that amyloid formation of p53 leads to loss of its specific DNA binding (Figure 1D).

Considering the loss of transcriptional activity of p53, we hypothesized that p53 amyloid formation may also lead to loss of key p53 functions such as triggering of apoptosis under stress conditions. To examine this, core fibril treated MCF 10A and HFF cells were subjected to Actinomycin D (ActD) treatment as a stress inducer (Figure 1E). Actinomycin D (ActD) treatment only to the untreated cells was used as a control. In both the cell lines, the core fibril treated cells showed resistance to apoptosis in the presence of stressor as compared to untreated cells (Figure 1E, S2A). This suggests the loss of apoptotic function of p53 due to amyloid formation.

Next, we hypothesized that similar to mutant p53 (Stein et al., 2019), p53 amyloids can also act as an oncoprotein. Previously, we had demonstrated that p53 amyloids can enhance the existing transformative properties of SH-SY5Y cells (neuroblastoma cell line) (Ghosh et al., 2017). We propose that similar changes might also induce the transformation of normal cells. Cells acquiring such transformative capabilities should exhibit various properties similar to transformed cells such as colony formation in soft agar, hyperproliferation and enhanced migration (Brosh and Rotter; Dong et al., 2013; Muller et al., 2011; Oren and Rotter, 2010). Indeed, core fibril treated MCF 10A and HFF cells display enhanced ability to form colonies in in the soft agar assay (cell transformation assay) as compared to the untreated controls (Figure 1F, S2B). This transformative potential is further assisted by their enhanced proliferation as evidenced by the higher number of cells expressing Ki67 (a cellular proliferation marker (Scholzen and Gerdes, 2000) as compared to untreated cells (Figure 1G, S2C). Further, the cell migration study, using wound healing assay (Liang et al., 2007) showed higher migration of core fibril treated MCF 10A cells (migration rate of ~13.61 μm/hr) compared to untreated cells (migration rate of ~3.44 μm /hr). A similar observation was also seen in HFF cells (Figure S2D, E). This property of enhanced migration, analogous to cancerous cells (Yamaguchi et al., 2005), indicates a gain of transformative properties by these cells.

### Mother-to-daughter cell transmission of p53 amyloid aggregates promotes cell survival

The role of p53 in pro-survival activities of a cell is tightly regulated and is integrated with its tumor suppressive function to prevent cancer induction (Reisman and Loging, 1998). Loss of regulation over these pathways provides the cells with a survival advantage. We aimed to understand whether the intracellular p53 amyloid aggregates can transmit from mother to daughter cells, which might also affect the survival or number of cell generations (passage number). To evaluate this, both the cells (MCF 10A and HFF) were treated with core fibrils and were passaged continuously for many generations. Untreated cells were used as control. Important to note that only the first generation of cells was treated with core fibril seeds. All subsequent generations did not receive any exogenous fibril treatment. Each cell generation after core fibril treatment was termed T1, T2 and so on (Figure S3A, B). Annexin-V PI assay coupled with FACS analysis of MCF 10A and HFF cells showed increased survival of core fibril treated cells (reduced apoptosis) over the generations as compared to untreated cells in corresponding passages (Figure 1H, S3C). At each passage, MCF 10A cells were immunostained using p53 DO-1 antibody for the detection of p53 in the cells. For core treated cells, p53 aggregates (predominantly punctate in the nucleus) were detected throughout all the cell generations (Figure 1I, S3D), indicating that p53 amyloids propagate through subsequent generations. Untreated cells did not show the presence of p53 aggregates (Figure S3D). Further, the presence of p53 amyloid fibrils was confirmed by dot blot assay using amyloid fibril specific OC antibody, in the later generations for both MCF 10A and HFF cells (Figure 1J, S3E). This suggests that p53 amyloid aggregates can be passed from one cell generation to the next, contributing to enhanced survival of the cells. Interestingly, the functional inactivation of p53 also persists through cell generations due to the propagation of p53 amyloids. This is evident as luciferase reporter assay using MCF 10A core fibril treated cells showed loss of p53 transcriptional activity both at T1 and T5 in contrast to untreated cells (Figure 2A). Interestingly, the loss of p53 transcriptional activity is more prominent at T5 in comparison to T1, which is consistent with the extent of p53 amyloid formation in cells (OC immunoreactivity, Figure 2A, 1J, S3E). This indicates that the amyloid form of p53 is no longer able to participate in cellular apoptotic mechanisms, allowing core fibril treated cells to overcome apoptosis for higher survival, whereas untreated cells undergo apoptosis in the continuous culture.

**Figure 2.**
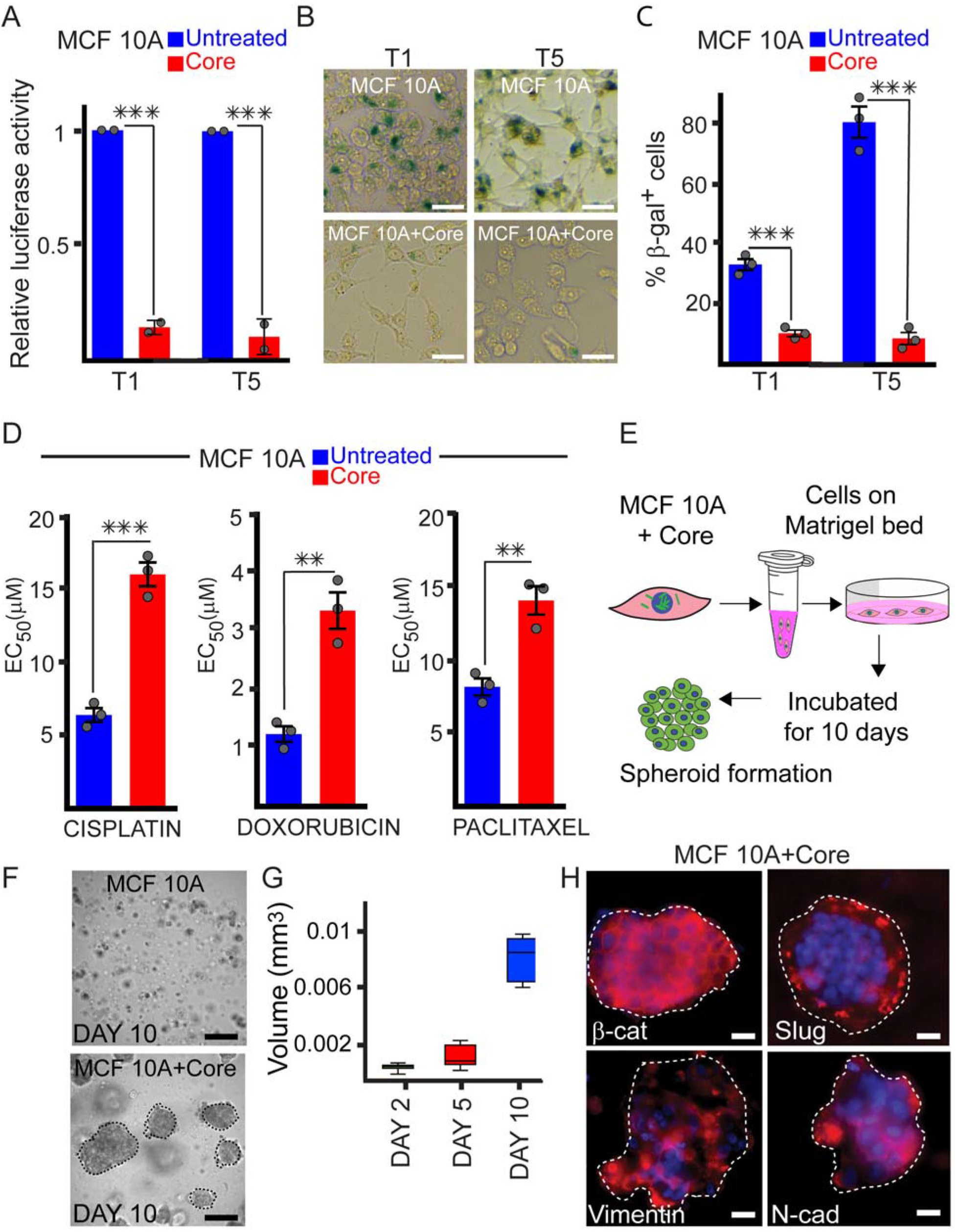
Loss of tumor suppressive and gain of oncogenic functions in fibril transformed cells. (A) Luciferase reporter assay showing loss of p53 transcriptional functionality at first generation (T1) and fifth generation (T5) after p53 fibril treatment. Untreated cells showing high luciferase activity indicating functionally active p53 in these cells. For (A), the values represent mean ± s.d, n=2 independent experiments. The p-value is 0.00064 at T1 and p-value 0.00373 at T5 with respect to the untreated control. (B) and (C) p53 amyloid formation confers resistance to cellular senescence. Senescence associated biomarker beta-galactosidase is quantified using X-gal. The core fibril treated cells showing the resistance to senescence over passages as compared to untreated cells. Scale bars are 40 μm. For (C), the values represent mean ± s.e.m, n=3 independent experiments. The p-value is 0.00045 at T1 and 0.002 at T5, calculated with respect to the untreated control. (D) Drug resistance in fibril treated cells using cisplatin, doxorubicin and paclitaxel. MTT assay quantifying cell viability in the presence of these drugs showing significantly higher EC_50_ values for the fibril treated cells as compared to untreated cells for all drugs. The values plotted represent mean ± s.e.m, n=3 independent experiments. The p values for cisplatin, doxorubicin and paclitaxel with respect to untreated cells are 0.00052, 0.00338 and 0.00631, respectively. (E) Schematic representation of experimental setup for the spheroid formation by p53 fibril transformed cells. The core fibril treated cells were passaged (till T5) and tested for spheroid formation in the presence of Matrigel. Untreated passaged cells (till T5) were used as control. (F) Spheroid formation by core fibril treated MCF 10A cell monitored for 10 days. Scale bars are 300 μm. (G) Spheroids formed by core fibrils treated cells showing an increase in spheroid volume over 10 days in 3D cell culture volume. (H) Immunofluorescence study of spheroids (day 10) showing expression of EMT markers, *β*-catenin (*β*-cat), Slug, Vimentin and N-cadherin (N-cad) by p53 core fibrils treated cells (at T5). Scale bars are 50 μm. The values plotted represent mean ± s.e.m, n=3 independent experiments. Statistical significance (***p ≤ 0.001, **p ≤ 0.01) for all experiments is determined by one-way ANOVA followed by Bonferroni multiple comparison post hoc test with 95% confidence interval.

### p53 amyloid formation confers cellular resistance against senescence and cancer drugs

p53 is known to regulate senescence and aging of cells through its tumor suppressive pathways (Itahana et al., 2001) and inactivation of p53 has been shown to induce reentry of cells into the cell cycle and rapid tumor progression (Kim et al., 2009). The p53 amyloid formation can, therefore, deregulate the pathways involved in activating senescence, thereby changing the fate of the cells. We examined senescence-associated beta-galactosidase (SA-*β*-gal) expression with chromogenic substrate X-Gal using a cytochemical assay (Gary and Kindell, 2005). Core fibril treated cells, both at early (T1) and later (T5) generation, showed a reduction in the percentage of SA-*β*-gal positive cells as compared to untreated control (at T1 and T5, respectively) (Figure 2B, C). This indicates that fibril treatment plays a role in the inactivation of senescence/aging pathways of cells, which further contributes to the cancer-like phenotype.

Furthermore, loss of p53 function has been established to play a critical role in conferring a high level of drug resistance to cancer cells against doxorubicin, cisplatin, alkylating agents, gemcitabine and microtubule targeting drugs (Hientz et al., 2017; Zhou et al., 2019a). p53 aggregation has also been shown to enhance platinum resistance in ovarian cancer cells (Yang-Hartwich et al., 2015). To examine the possible drug resistance of cells due to p53 amyloid formation, the toxicity of fibril transformed cells (at T5) were analyzed using MTT assay in presence and absence of cisplatin, doxorubicin and paclitaxel, which are well-known drugs used against breast cancer (Onda et al., 2004). Core fibril treated cells showed significantly higher EC_50_ values for all these drugs as compared to untreated control (Figure 2D). The survival trend of untreated cells and core fibril treated cells was plotted for each of the drug concentrations (Figure S3F). This validates that p53 amyloid formation can induce drug resistance in cells, which can aid in cancer development and therapeutic resistance.

### p53 amyloid formation induces tumorigenic properties in cells

As MCF 10A cells acquire a transformative phenotype after p53 amyloid formation, they might also show gain-of-oncogenic traits, characteristic of tumor cells, such as the formation of spheroids in three-dimensional culture (Tevis et al., 2017). To examine this, we used Matrigel mediated 3D cell culture method (Figure 2E). Core fibril treated MCF 10A cells at T5 passage showed 3D spheroid formation with an average diameter of ~100 μm on day 2 (Figure 2F, S3G, H). The volume of spheroids was calculated from the diameter and was found to increase gradually over 10 days (Figure 2G), indicating proliferation and migration of core fibril treated cells. Untreated cells, however, did not show significant cellular aggregation to form distinct 3D spheroids (Figure 2F). Further, spheroids showed positive staining with Calcein-AM dye indicating their viability (Figure S3I). As p53 maintains a transcriptional program to prevent epithelial to mesenchymal transition (EMT) of cells (Chang et al., 2011), the loss of this p53 activity may induce EMT-like phenotype in cells (Muller et al., 2011). In this context, the amyloid-induced 3D spheroids on day 10 showed high expression of EMT markers like *β*-catenin, Slug, Vimentin and N-cadherin as evident by immunofluorescence (Figure 2H). This suggests that cells with p53 amyloid aggregates promote EMT potential, which is known to contribute to cancer progression (Derynck and Weinberg, 2019; Roche, 2018). Finally, to confirm the p53 amyloid-mediated gain of tumorigenicity by the cells, core fibrils treated cells at T5 were injected in SCID mice to examine tumor induction in a mouse xenograft model (Figure 3A). The untreated MCF 10A (at T5) and MCF 7 cells were also injected as negative and positive controls, respectively. The injection of fibril treated cells in the mammary fat pad of female mice showed gradual tumor generation after 2 weeks while no tumor was observed for untreated MCF 10A cells (Figure 3B). MCF 7 cells, the positive control for the xenograft also showed tumor development in the animals. The increase in tumor volume was calculated up to 5 weeks, after which the animals were sacrificed and tumors were isolated (Figure 3C, D). Further, the PET/CT scan was used as a non-invasive technique to visualize the tumor anatomy and assay the metabolic activity of the tumor (Mulero et al., 2011). The tumors were first identified with small animal CT followed by a small animal PET scan for the evaluation of ^18^F-FDG accumulation. The tumors showed an accumulation of ^18^F-FDG, indicating active metabolic activity (Figure 3E). Increased standardized uptake values (SUVs) of ^18^F-FDG values for MCF 7 injected animals (1366.7 ± 225.6) and core fibril treated MCF 10A injected animals (1051.6 ± 145.5) reflect the viability and metabolic activity of cells in the tumor (Figure 3F). The histology of excised tissue (tumor/normal) was examined using Hematoxylin and Eosin staining (Figure 3G). Tumor tissue from animals injected with core fibril treated MCF 10A showed loss of normal ductal-lobular architecture with a high degree of nuclear pleomorphism. The extensive nuclear staining by hematoxylin indicated hyperproliferative cells in the tumors with loss of normal tissue histology, similar to MCF 7 tumor (Figure 3G). In contrast, normal tissue morphology was observed in untreated MCF 10A injected animals (Figure S3J).

**Figure 3.**
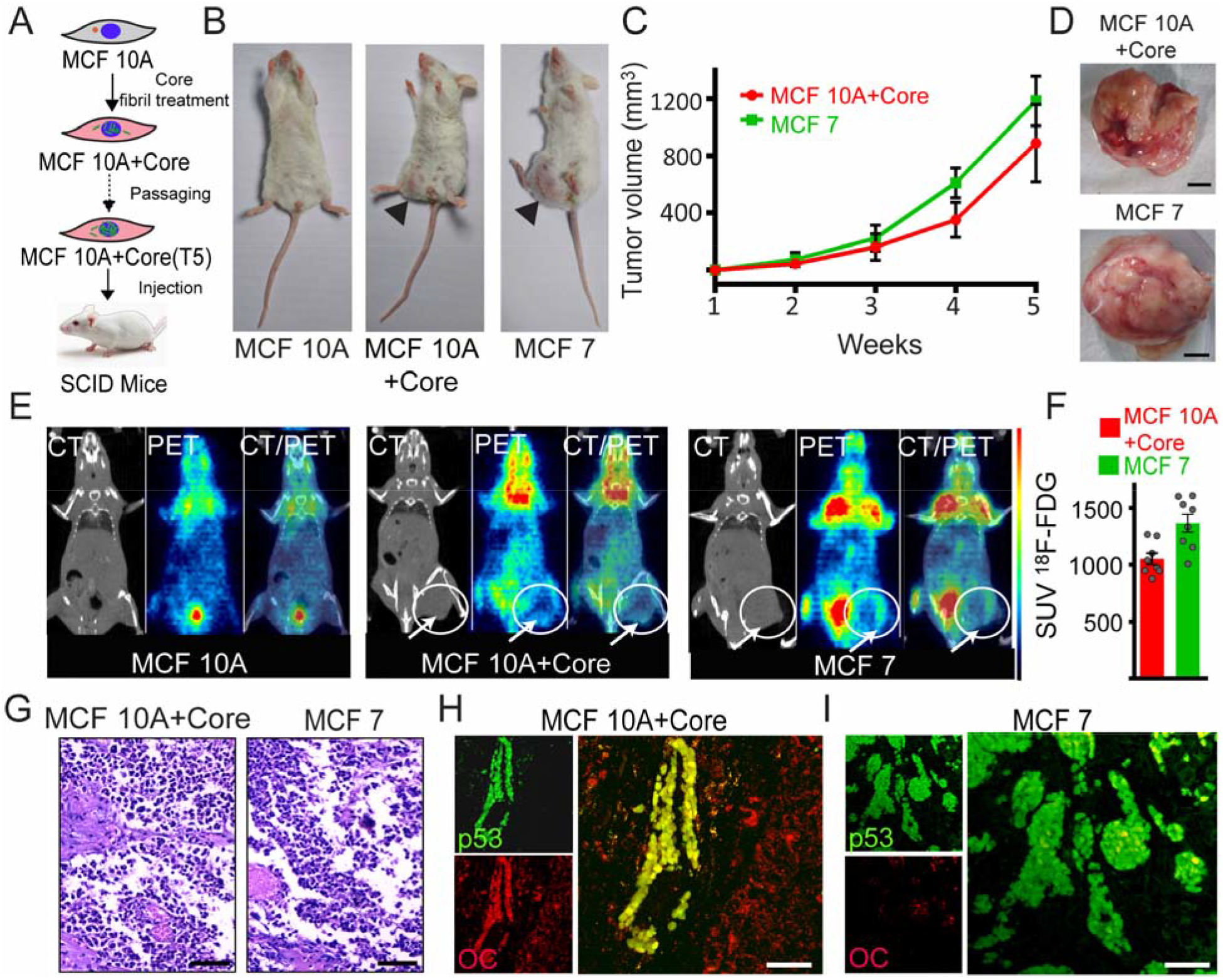
p53 amyloid formation induces tumorigenesis in the mouse xenograft model. (A) Schematic showing injection of MCF 10A core fibril treated cells in the SCID mice for the establishment of xenograft. (B) Tumor formation by MCF 10A core fibril treated cells (Core) similar to MCF 7 (positive control) when injected in the mammary fat pad of SCID mice. Both sets of mice showed the appearance of tumors in 2-3 weeks. The mice with an injection of only MCF 10A cells did not show any tumor induction. (C) Progression of tumor volume over time showing tumor growth of MCF 7 tumors and core fibrils treated MCF 10A tumors. (D) Image of excised MCF 7 tumor and core fibrils treated MCF 10A tumor from mouse xenografts. Scale bars are 20 mm. (E) CT and PET scanning to detect tumors in mice. Tumor-bearing animals imaged using PET/CT scan showing the anatomy of the tumors by CT and metabolic activity as measured by uptake of radiolabeled ^18^F-FDG by PET. PET data showing the active tumor formation by fibril treated MCF 10A cells similar to MCF 7 cells (ROI marked with arrows). (F) Quantification of ^18^F-FDG uptake values from PET scan of fibril treated MCF10A injected and MCF 7 cells injected tumors from the region of interest. Increased ^18^F-FDG uptake indicates the higher metabolic activity of cells in the tumor. Values plotted represent mean ± s.e.m, n=8 animals. (G) H & E staining showing differential staining with hematoxylin and eosin for tumor tissues generated by injecting p53 core fibril treated MCF 10A cells and MCF 7 cells. The tissues showing intense hematoxylin staining indicating the hyper-proliferation of cells in the tumor produced by MCF 10A core cells similar to MCF 7 cells. Scale bars are 100 μm. (H) and (I) Immunohistochemistry of tumor tissue sectioned from p53 core fibril treated MCF 10A cell tumor (H) and MCF 7 (I) cell-derived tumor using p53 DO-1 antibody and amyloid specific OC antibody. The tumor tissue from p53 core fibrils treated MCF 10A cells showed colocalization of p53 and amyloid staining indicating the presence of p53 amyloids in the proliferating tumor tissue. MCF 7 tumor showed p53 stabilization without colocalization with OC indicating the absence of p53 amyloid aggregates. Scale bars are 50 μm.

The immunohistochemistry of tumor tissues using p53 DO-1 antibody and amyloid specific antibody (OC) revealed the presence of p53 aggregates in the tumor tissue (Figure 3H) and negligible p53 expression was observed in control tissue from MCF 10A only injected animals (Figure S3J). Tumors induced by MCF 7 cells showed stabilized expression of p53 but without colocalization with OC, indicating the absence of p53 amyloid aggregates (Figure 3I). These results confirmed the ability of p53 amyloid aggregates to impart tumorigenic properties to normal cells (MCF 10A).

### Gene expression changes due to p53 amyloid formation

We performed microarray analysis to understand the distinct cellular pathways modulated by p53 amyloid formation and how these pathways contribute to cellular transformation leading to cancer initiation. To understand the immediate effect of p53 amyloid formation and its eventual consequences, we performed global gene expression analysis of p53 core fibril treated MCF 10A cells at two different generations-initial (T1) and later (T5) generation (Figure 4A). Bioinformatic analysis revealed 2160 transcripts (at T1) and 1008 transcripts (at T5) were differentially regulated upon core fibril treatment as compared to the untreated control. 425 genes were seen to overlap between T1 (19% of 2484 transcripts) and T5 (42% of 1112 transcripts) highlighting the central role of p53 in regulating gene expression (Figure 4B, C). The higher number of genes influenced at the T1, compared to T5, suggests that p53 amyloid formation in cells deregulates many cellular pathways at the beginning, contributing to the survival of cells. We performed a pathway enrichment analysis to examine the cellular response upon loss of p53 exclusively due to p53 aggregation, using Metascape (Zhou et al., 2019b). Significantly affected genes in both T1 and T5 were annotated in terms of biological processes, cellular components and molecular function (Figure 4D).

**Figure 4.**
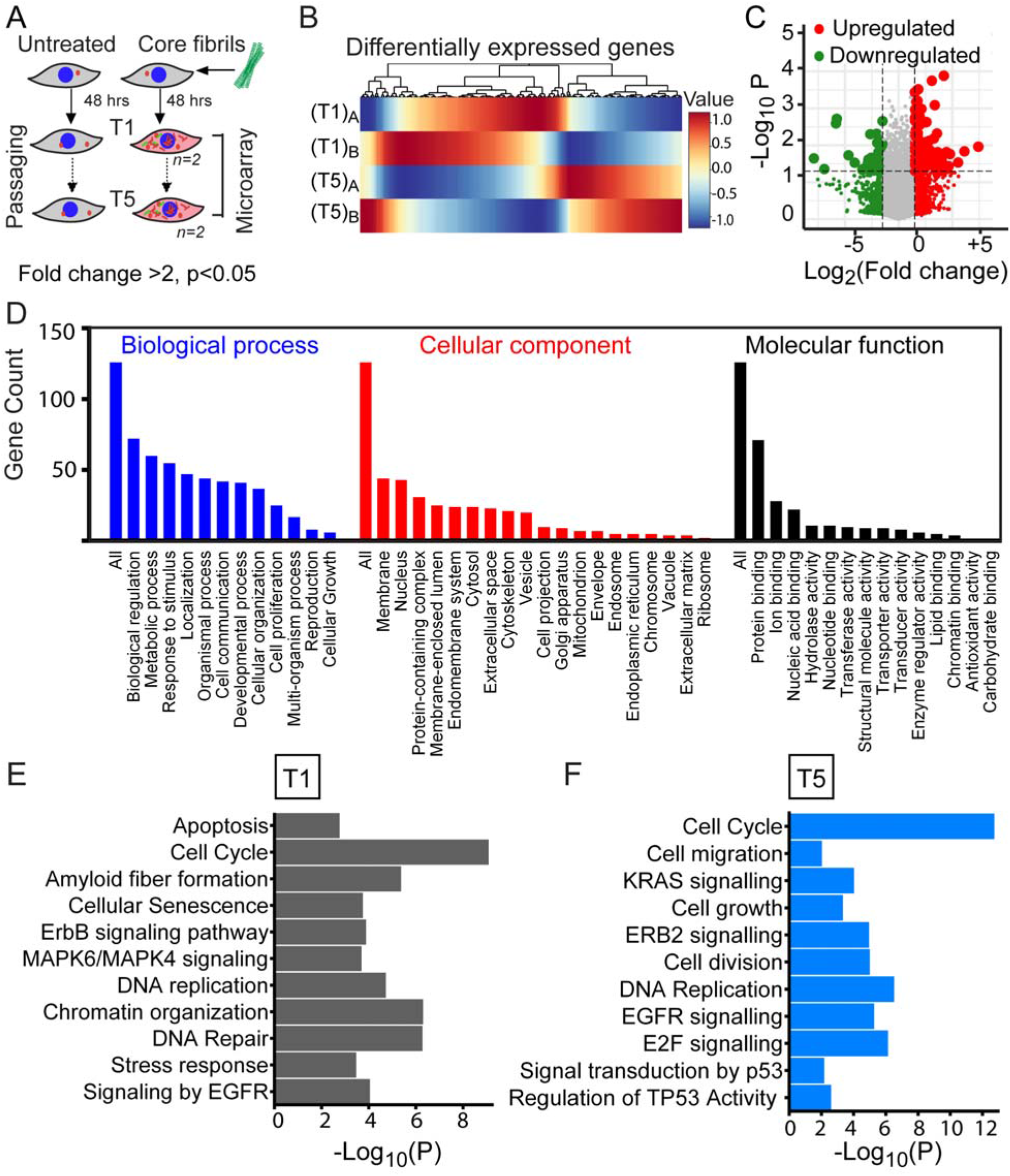
Gene expression analysis by microarray showing pathways affected by p53 amyloid formation. (A) Schematic representation of cells that are used for microarray analysis. (B) Heatmap of genes that are differentially regulated upon core fibril treatment in MCF 10A cells (for two different passages, T1 and T5). (T1)_A_ and (T1)_B_ are two biological replicates for the T1 passage. Similarly, two biological replicates are done for cells in T5 passages. (C) Volcano plot representing the log_2_ fold change plotted against the −log_10_ P-value. Differentially expressed genes are represented in large size dots of green (significantly upregulated) and red (significantly downregulated) colour. (D) Gene functions of significantly affected genes annotated in terms of the molecular function of genes, biological processes and the cellular components are shown. (E) and (F) Metascape analysis showing pathways that are significantly affected by p53 amyloid formation at T1 and T5 generations, respectively. At T1 generation, cell cycle, apoptosis and DNA repair are significantly perturbed. The T5 generation showing perturbation of pathways involved in cell cycle, DNA replication, cell migration and cell division.

At T1 passage, downregulation of Cdc6, CDK4, Cdc45, PCNA, E2F2, Rad51 and TIMELESS genes involved in cell cycle checkpoints, senescence and DNA repair processes, respectively, was observed, indicative of the loss of p53 tumor suppressive network, suggesting loss of native p53 regulatory control at an initial stage (Figure 4E). The pro-apoptotic genes (DAPK3) and genes involved in proofreading and repair during DNA replication (TREX1 and EXO1) were also downregulated (Figure S4A). In contrast, anti-apoptotic genes and genes involved in chromatin remodeling (AKT1, SEPT4, BAD, KDM4B and HDAC1) were upregulated. Further, the data showed upregulation of genes, ERBB2 (Her2), MAP2K1, EGF, involved in promoting cell growth, signaling and proliferation (Figure S4A). The data indicate that p53 amyloid formation at an early passage (T1) caused an immediate and drastic effect on tumor suppressive pathways of the cell. At the later stage (T5), the regulatory genes associated with cell cycle and cell division (such as CdcA2, Cdc45) showed decreased expression; whereas genes having a role in cellular architecture, migration and growth (such as Vimentin, PECAM, WNT11 and STAT3) showed enhanced expression (Figure 4F). Kras and multiple protein kinase signaling pathway genes involved in many cancers (Brognard and Hunter, 2011; Martin, 2003), were also upregulated in our datasets (Figure S4B). Hence, at the initial stage (T1), p53 loss of function is more drastic and at a later stage, the gain-of-oncogenic function pathways are seen to be significantly altered, which is responsible for the tumorigenic nature of the T5 stage of cells.

Moreover, from the entire dataset of genes, we specifically mapped expression levels for a subset of genes (n=76) which are a crucial part of the p53 signaling network (Joerger and Fersht, 2016; Soragni et al., 2016). These direct targets of p53 (such as p63, p73, CCND1, NOXA1, GADD45 and PCNA) showed deregulation at both T1 and T5 generation (Figure S4C). This suggests that p53 amyloid formation also affects the downstream regulatory function. Further, to evaluate how the expression of p53 target genes can gradually change across the cell generations (with a simultaneous increase in p53 aggregates in the cells), we evaluated selective p53-linked genes for their relative expression from T1 to T5 passages using quantitative PCR (qRT-PCR) across cell generations (Figure S4D). The direct p53 target p21 showed decreased expression through the passages. However, pro-cancer genes like CCND2, MAPK showed a gradual increase in expression and anti-cancer genes like BAX and DDB2 showed concomitant down-regulation (Figure S4D). Altogether, this data suggests that, mainly, pathways of cell cycle, apoptosis and pro-proliferative signaling are majorly affected, leading to a gradual gain of oncogenic phenotype in core fibril treated cells.

### Proteome-wide changes in cells due to p53 amyloid formation

We further examined the proteome of fibril treated cells to elucidate changes in the cellular proteome and protein-protein interactions due to p53 amyloids using iTRAQ labeling coupled with LC-MS/MS analysis (Datta et al., 2017; Zhao et al., 2016). Similar to the microarray analysis, we profiled the p53 core fibril treated MCF 10A cells at two different time points, initial (T1) and later (T5) generations. Expression levels of total 412 at T1 and 261 at T5 proteins were found to be altered due to core fibril treatment compared to the untreated control (Figure 5A, B). We further mapped the altered proteins to their functional classes in terms of cellular localization and molecular function (Figure S5A, B) using STRING (version 11.0) (Szklarczyk et al., 2018). At T1 passage, proteins involved in the regulation of cell cycle (such as CDK1, CCAR1), apoptosis (such as Bcl2, PDCD6), p53-dependent DNA damage checkpoints decreased significantly; whereas proteins involved in MAPK4 signaling (MAP3K10) and Wnt signaling showed higher expression levels (Figure 5C, S5C). Apart from these processes, chaperones belonging to Hsp70 and Hsp90 families showed significantly increased levels (Figure 5C). At the later generation (T5), cell cycle regulatory proteins like CDC42 and protein associated with cell death showed decreased levels. In contrast, the proteins implicated in translational machinery (such as EIF4, EFF2) and cell proliferation pathways (such as CDK2, CRNKL1, MAPK1) showed higher levels of expression (Figure S5D, E). Interestingly, at both T1 and T5, proteins in pathways governing ubiquitination, unfolded protein response and proteasomal degradation machinery (UBP1, PRS 10, PSA3) were upregulated due to presence of p53 aggregates in the cells (Figure 5C, S5E). Such changes were also demonstrated previously when cells were treated with artificial *β*-sheet proteins that were designed to form amyloid-like fibrils (Olzscha et al., 2011). This indicates that apart from p53 specific pathways, p53 amyloid formation also causes aberrant changes in protein quality control and clearance mechanisms, possibly affecting multiple cellular functions. In addition to this, we found that at both T1 and T5 passages, proteins involved in metabolic processes such as glycolysis, citric acid cycle and amino acid synthesis showed increased levels of expression (Figure 5C, S5E). Cancer cells are known to rewire metabolic pathways and display higher metabolic rates which provide them with a growth advantage over normal cells (Hsu and Sabatini, 2008). These upregulated metabolic proteins suggest that carbohydrate and protein metabolism in fibril treated cells is significantly altered, contributing to the proliferation/survival.

**Figure 5.**
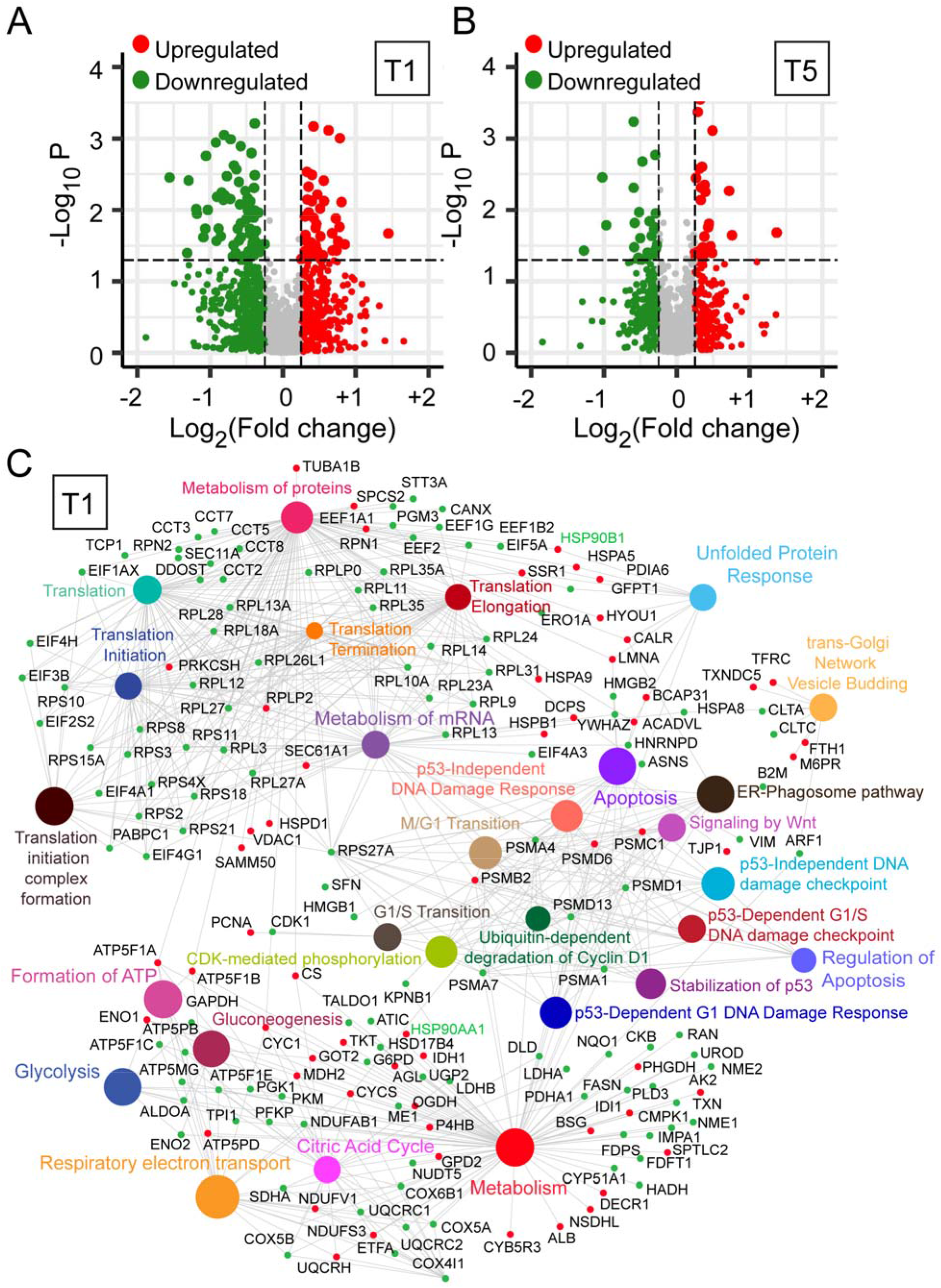
The changes in proteome due to p53 amyloid formation in cells. Core fibril treated cells were used for proteomic analysis at T1 and T5 passages and the protein levels are compared to the untreated cells. Duplicates of each passage were used for analysis. (A) and (B) Volcano plot displaying log_10_ (p-value) vs log_2_ fold-change values corresponding to differentially expressed proteins at T1 passage (A) and T5 passage (B). The red dots represent the overexpressed proteins and the green dots representing the proteins with lowered expression compared to the untreated control cells. (C) Network analysis of proteins, which are significantly altered upon fibril treatment to the cells was performed using “NetworkAnalyst 3.0”. Bubble plot representing proteins as nodes and the clusters of proteins as dense circles in particular pathways are shown. At T1, the significant fold changes were observed in proteins that belong to the pathways regulating apoptosis, cell cycle and senescence of cells. Further, the proteins involved in the carbohydrate metabolism also showed a higher level of expression, which consequently modify the pathways of glucose utilization like glycolysis and citric acid cycle (as shown). These alterations in protein levels contribute to the loss of p53-mediated survival regulation and parallelly evolve robust survival traits in fibril treated cells.

Thus, overall, global analysis of gene expression and proteome-wide changes suggest that cells with p53 amyloids show a loss of function in terms of apoptotic and cell cycle arrest pathways, thus conferring a growth advantage to p53 amyloid containing cells. A concomitant gain of oncogenic functions via increased proliferative signaling and a higher rate of metabolism rewires the cellular pathways and leads the cells to a cancer-like phenotype.

### Rescue of p53 aggregation reverses the cancer-like traits in cells and resists tumorigenesis

If the observed transformed phenotype of the cells is initiated due to core fibrils treatment (i.e., endogenous p53 aggregation), then rescuing the p53 aggregation using inhibitor should reverse the oncogenic changes. In this context, Soragni *et al* have developed a peptide inhibitor based on aggregation-prone region of p53 termed as rescue peptide (RP) (Soragni et al., 2016). We synthesized this peptide and exogenously added to the cells, which possess the aggregated p53 (at T5). Immunofluorescence staining of p53 in cells showed reduced p53 aggregates in the nucleus as compared to the untreated control cells (Figure 6A). We further checked if the transcriptional function of p53 is restored after RP treatment. Indeed, p53 disaggregation led to the rescue of p53 sequence-specific binding and transcriptional function, as demonstrated by higher luciferase activity compared to untreated cells (−RP) (Figure 6B). Further, to check if the apoptotic function of p53 is restored due to p53 disaggregation by inhibitor peptide, Annexin-V PI assay followed by FACS analysis was done. RP treated cells showed loss of viability with an increase in apoptosis as seen in FACS data (Figure 6C, S6A). Since these cells are at T5 generation where ideally the MCF 10A cells without p53 aggregation (untreated) do not maintain their viability (Figure 1H), high apoptotic death is observed even in absence of external stressor (due to RP treatment). Moreover, the RP treated cells showed reduced migratory potential (Figure 6D, S6B) and decreased colony formation potential in soft agar assay (Figure 6E, S6C), comparable to untreated MCF 10A cells without p53 aggregation.

**Figure 6.**
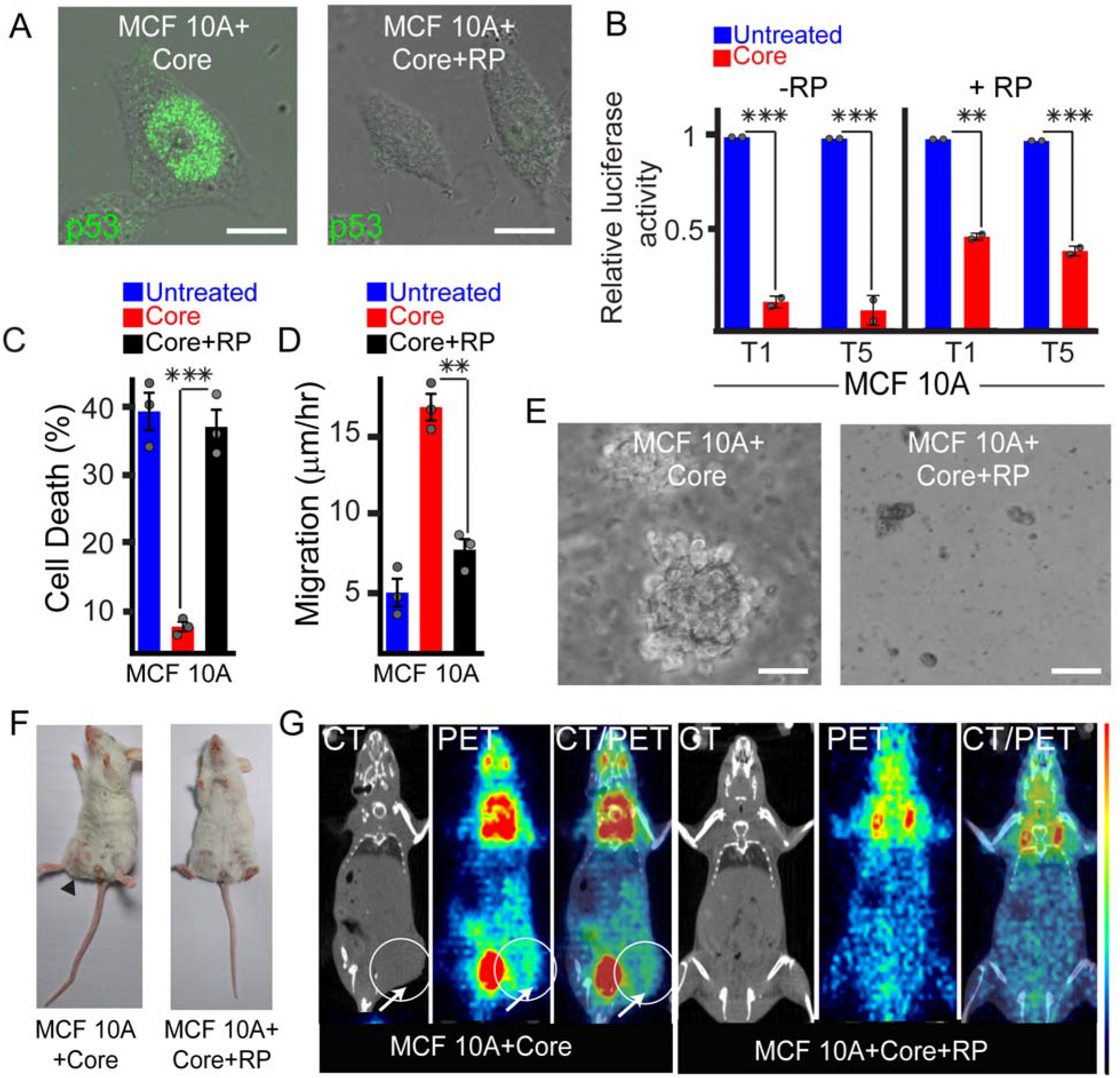
Dis-aggregation of p53 amyloid formation rescues the native p53 function. (A) Core fibrils treated cells (at T5) when incubated with 10 μM rescue peptide (RP) for 16 hrs showing the disappearance of p53 punctate, indicating disaggregation of p53. Scale bars are 10 μm. (B) Relative luciferase activity of fibril treated cells showing a significant increase after incubation with rescue peptide, suggesting that disaggregated p53 regains its native DNA binding property and transcriptional activity. Untreated cells showing a higher luciferase activity, indicating functionally active p53 in these cells. The p values for −RP cells at T1 and T5 with respect to untreated cells are 0.00064 and 0.00373, respectively. The p values for +RP cells at T1 and T5 with respect to untreated cells are 0.001673 and 0.0021, respectively. (C) Annexin-V PI assay followed by quantification by FACS analysis showing increased apoptosis of core fibrils treated cells when incubated with RP. Values represent mean ± s.e.m, n=3 independent experiments. the p-value for Core+RP cells is calculated to be 0.00037 with respect to core only cells. (D) Cell motility using scratch assay showing the reduction of migration rate by rescue peptide treated cells compared to only core fibril transformed cells. (E) Soft agar assay showing no colony formation by rescue peptide treated cells as compared to core fibril treated cells, which showed colony formation. Scale bars are 50 μm. (F) Effect of rescue peptide (RP) on p53 core fibrils treated cells for its tumor induction in the mouse xenograft model. Injection of core fibril treated MCF 10A cells followed by RP treatment showing no tumor formation; while tumor formation was observed with RP untreated cells (core fibril treated at T5). This suggests that RP treatment and consequent p53 disaggregation reduces the tumorigenic potential of the core fibrils treated MCF 10A cells. (G) CT/PET scan showing no tumor formation by RP treated MCF 10A cells (core fibrils treated at T5), while the same cells without RP treatment showed metabolically active tumor (with increased ^18^F-FDG uptake). This validated the reduction of the tumorigenic potential of cells due to p53 disaggregation by RP treatment. Statistical significance (***p ≤ 0.001, **p ≤ 0.01) is determined by one-way ANOVA followed by Bonferroni multiple comparison post hoc test with 95% confidence interval.

Further, when the rescue peptide treated cells were injected in mice, these cells did not induce tumor formation as compared to fibril-transformed cells (Figure S6D, 6F). This suggests that rescue peptide treatment partially reverts the cellular status similar to untreated MCF 10A cells. Core fibril treated MCF 10A cells xenograft tumor showed neoplastic cells with extensive hematoxylin staining of the nuclei, indicating the presence of hyperproliferative cells. However, rescue peptide treated cells neither showed tumor formation in mice nor hyperproliferative cells in the tissue of the injected area (Figure S6E). CT imaging further confirmed the anatomy of tumor in core fibril treated MCF 10A injected animals with increased ^18^F-FDG uptake; whereas no signal was detected for animals injected with the same cells but treated with RP (Figure 6G). This suggests that RP treatment results in p53 disaggregation and cells are unable to form tumor xenograft in mice.

## Discussion

Protein aggregation and amyloid formation are linked with many human neurodegenerative diseases, such as Alzheimer’s, Parkinson’s, and prion disease (Dobson, 2002; Dobson, 2001, 2003; Soto, 2003). In these diseases, the cellular toxicity is associated with (i) loss of protein native function and/or (ii) gain of toxic function as protein aggregate serves as a toxic entity (Eisenberg and Jucker, 2012; Winklhofer et al., 2008). Moreover, many amyloids are known to be associated with native functions of the host organism suggesting that amyloid fibrils, in general, might not always be toxic (Maji et al., 2009; Maury, 2009). The most recent addition to the amyloid family of diseases is p53, implicated in cancer pathogenesis, wherein p53 loss of tumor suppressor function is suggested to be associated with its amyloid formation (Costa et al., 2016; Ghosh et al., 2017; Lasagna-Reeves et al., 2013). The recent demonstration of p53 fibril uptake by cells (Forget et al., 2013) and cell-to-cell transmission of p53 aggregates (Ghosh et al., 2017) raises the possibility that p53 amyloids can act as a transmissible prion protein. It was hypothesized that amyloidogenic aggregates of p53 can, thus, spread loss of function, dominant-negative and gain-of-function phenotypes from cell-to-cell (Navalkar et al., 2020). Therefore, it becomes essential to understand the molecular mechanisms associated with p53 amyloid formation in the cells, which may lead to prolonged survival of cells and cancer initiation.

To study this, here, we evaluate if wild-type p53 in normal cells, induced to form amyloid aggregates due to the exogenous addition of amyloid seeds, can cause the normal cells to undergo transformation, acquiring cancerous phenotype. p53 aggregation was induced in MCF 10A and HFF cells (mostly localized in the nucleus), which led to loss of p53 transcriptional function (Figure 1B, S1E, 1D). In this context, previous studies have shown that cytoplasmic p53 amyloid formation not only inhibits p53 translocation into the nucleus but also sequesters other proteins including other tumor suppressors and p53 paralogs (p63, p73), interfering with their native functions (Xu et al., 2011). Furthermore, when p53 amyloids are formed in the nucleus, it will not able to bind to the cognate DNA sequence for transcriptional activities, rather, it might bind non-specifically to alternative sequence elements and initiate transcription of genes for malfunction, as observed for mutant p53 (Göhler et al., 2005; Muller and Vousden, 2014). Indeed, MCF 10A and HFF cells containing p53 amyloids showed resistance to apoptosis, when exposed to stress conditions and also introduced pro-metastatic properties in cells, (such as enhanced motility and transformation potential), resembling cancer cells (Figure 1E, 1F, S2D).

p53 amyloids can be transmitted from one generation to next generation in cells (mother to daughter cells) leading to an increase in survival in terms of the number of generations (increase passage number), all of which retain the p53 amyloid aggregates. The presence of p53 amyloids also induced hyperproliferation of cells, decreased senescence and showed increased resistance against cytotoxic drugs (Figure 1G, H, 2B-F). These cellular properties (similar to mutant p53 (Mantovani et al., 2019; Oren and Rotter, 2010)) might cause interference in a multitude of cellular pathways, which are likely to contribute to oncogenic properties or transformation of cells. This cellular transformation due to p53 aggregation is further evident by the fact that the cells containing p53 aggregates display the ability to form 3D spheroids (in contrast to cells with no p53 aggregation) and exhibit the expression of proteins involved in EMT (Figure 2H-J). The direct evidence that p53 aggregates confer tumor-like properties to the cells is the observation that cells with p53 amyloid aggregates induce tumorigenesis in mice (Figure 3). Further, the tumor tissue demonstrated the presence of p53 aggregates, establishing a direct link between p53 amyloid formation and tumorigenesis. In this regard, the accumulation and subsequent aggregation of p53 has also been observed in clinical biopsy tissues of neuroblastoma, retinoblastoma, breast and colon cancers (De Smet et al., 2017; Levy et al., 2011; Moll et al., 1995). Understanding the consequences of p53 aggregation in relation to cancer progression, therefore, promises to be an important undertaking. Based on our microarray and proteomics dataset, we present here a mechanistic model for demonstrating the consequences of p53 amyloid formation in terms of cellular pathways, which are disrupted (Figure S7). Essentially, p53 amyloid formation can lead to the downregulation of the tumor suppressive and regulatory pathways at an early stage and gradually introduce the gain of oncogenic function via the upregulation of the cell cycle and cell proliferation pathways (Figure 4, 5). Our study proves that p53 amyloids can uniquely trigger oncogenic pathways with prolonged expression of oncogenes, which can accumulate over cell generations, conferring the tumorigenic nature to the cells. Importantly, disaggregation p53 partially restores the native function of p53 (Figure 6) and we observe that rescued p53 behaves similarly to the wild-type folded form, ultimately triggering cell death. Dis-aggregation of p53 diminishes the tumorigenic propensity induced by exogenous addition of fibril seeds. Our study, thus, establishes cancer as a protein misfolding disease and demonstrates that disaggregation of misfolded p53 can be used as an alternative approach for targeting a specific subset of cancers, which initiate due to p53 amyloid formation.

## Accession Numbers

The NCBI Gene Expression Omnibus accession number to download the microarray dataset is GEO: GSE150522. The mass spectrometry proteomics data has been deposited to the ProteomeXchange Consortium via PRIDE partner repository with the dataset identifier PXD019498.

## Statistical Analysis

The statistical significance was calculated by one-way ANOVA followed by Bonferroni Multiple Comparison post hoc test. The p-value for the significance is *p < 0.05, **p < 0.01, ***p < 0.001; non-significant (NS p > 0.05). KaleidaGraph, version 4.1 software was used for calculating the statistical significance. No outliers were excluded from analysis and no assessment was made for outliers and normality of data.

## Author contributions

S.K.M and A.N conceived the project and designed the experiments. A.N carried out all the experiments unless stated otherwise. S.P performed the senescence assay, reporter assay along with sample preparation and analysis of microarray. N.S carried out the spheroid model-based experiments. A.K and S.S acquired the proteomics data and helped in analysis. K.P performed confocal imaging. B.M helped in performing the CT/PET scan of the animals. S.K.M, A.N and S.P wrote the manuscript. All authors have proof-read and approved the manuscript.

## Acknowledgments

We acknowledge IIT Bombay central facilities for FACS and confocal microscopy for performing the experiments. We also acknowledge Genotypic Technology Private Limited Bangalore for microarray processing and verifying the data analysis reported in this publication. We are thankful to Prof. Venkatesh for his help in the analysis of microarray data. We are thankful to Prof. Charles Glabe, UC Irvine, USA for the kind gift of OC antibody. The authors acknowledge Lady Tata Memorial Trust and DST-SERB (EMR/2014/001233), Government of India for financial support. Authors acknowledge Dr. Shinjinee Sengupta, Dr. Nitu Singh and Soumik Ray for their inputs on the manuscript. A.N acknowledges UGC-CSIR, Government of India for the fellowship. S.P acknowledges DST-SERB, Government of India for funding and fellowship.

## Conflict of interest

The authors declare no conflict of interest.

## Supplementary information

Supplementary information includes experimental methods, seven figures, and one table.

## Notes

### Competing Interest Statement

The authors have declared no competing interest.

