## Supplementary Information for "Induction of intracellular wild-type p53 amyloids leading to cellular transformation and tumor formation in mice"

**Experimental Methods**

**Supplementary Figures**

**Supplementary Table**

**Supplementary References**

**Experimental Methods**

**Chemicals and reagents**

All reagents and chemicals used for the studies were of the highest quality and were purchased either from Merck (Darmstadt, Germany) or Sigma-Aldrich (St. Louis, MO, USA). Double-distilled, de-ionized water was obtained using a Milli-Q system (Millipore Corp., Bedford, MA, USA) and autoclaved before use. Rescue peptide was custom synthesized by USV Limited, Mumbai, India by solid-state peptide synthesis method with >95% purity, peptide mass and purity was confirmed using HPLC and MALDI-TOF mass spectrometry.

**Protein expression and purification**

p53 core protein was expressed and purified according to our previously published protocol (Ghosh et al., 2017) using p53 core domain plasmid (Addgene #24866). Briefly*,* p53 core domain plasmid was transformed in *E. coli* BL21 (DE3) to express and purify p53 core protein with N-terminal His6 tag. 1 mM isopropyl β-D-1- thiogalactopyranoside (IPTG, Himedia, India) was used to induce p53 core expression at 25 °C for 8 hrs. After induction, the cells were harvested and lysed in 50 mM sodium phosphate, pH 8.0 containing 0.3 M NaCl and 1 mM PMSF (Sigma Aldrich, USA). The cell pellet was sonicated at 40% amplitude for 15 mins with 3 seconds on and 3 seconds off cycle. The cell lysate was centrifuged at 8000 r.p.m at 4 °C for 45 mins. The supernatant was collected and added to Ni^2+^ sepharose affinity chromatography column to purify His-tagged p53 core protein from the cell lysate. The elution of the protein was done using the imidazole gradient from 100-250 mM. The protein containing solution was then dialyzed against 50 mM sodium phosphate buffer containing 0.3 M NaCl, pH 7.4, to remove the imidazole. The purity of protein fractions was confirmed using 12% SDS–polyacrylamide gel electrophoresis (SDS–PAGE) and core monomer protein concentration was measured using UV absorption considering molar absorptivity of protein (ɛ) as 17420 M^-1^.cm^-1^

**Aggregation of p53 core domain *in vitro***

p53 core monomer protein at a concentration of ~50 μM (500 μl) in 50 mM sodium phosphate buffer (pH 7.4, containing 0.3 M NaCl, 0.01% sodium azide) and equimolar concentration of chondroitin sulphate A (CSA, Sigma Aldrich, USA) dissolved in the same buffer were mixed. The protein solution was then used for setting up *in vitro* aggregation of p53 core domain according to the previously published protocol (Ghosh et al., 2017). Briefly, the solution was incubated at 37 °C for 96 hrs with slight agitation. After incubation, the protein solution was centrifuged at 45,000 r.p.m. for 45 min at 4 °C. The fibril pellet was collected and was resuspended in PBS, pH 7.4.

**Circular dichroism (CD) spectroscopy**

Circular dichroism spectroscopy of the p53 core monomeric protein (0 hrs) and p53 core amyloid fibrils (96 hrs) was done to confirm the structural transition and amyloid formation of p53 core. For this, 200 μl p53 core monomeric protein or resuspended p53 core fibrils (10 μΜ) were taken in washed quartz cell (Hellma, Forest Hills, NY) with a path length of 0.1 cm. CD spectra were obtained by CD instrument (JASCO-1500) between the wavelength range of 200-260 nm at room temperature. For each sample, 3 accumulations were acquired and the average was taken. Smoothing and buffer subtraction was done for the processing of raw data.

**Thioflavin T (ThT) fluorescence**

To evaluate the amyloid nature of p53 core fibrils, the ThioflavinT binding was measured for p53 core monomeric protein (0 hrs) and p53 core amyloid fibrils (96 hrs). For this, 200 μl p53 core monomeric protein or resuspended p53 core fibrils (10 μΜ) were used and 2 μl of 1 mM of ThT prepared in Tris-HCl buffer (pH 8.0, containing 0.01% sodium azide) was added. ThT fluorescence was measured with an excitation wavelength of 450 nm and an emission range of 460-500 nm (slit width of 5 nm) using Horiba-Jobin Yvon (Fluomax4) instrument. Fluorescence intensity at 480 nm for all the samples was noted for ThT data analysis. Three independent sets of experiments were done for each of the samples.

**Electron microscopy and fibril seed preparation**

The fibril morphology of the aggregated p53 core domain was confirmed using Transmission electron microscopy (TEM). After p53 aggregation *in vitro*, the protein solution was centrifuged at 45,000 r.p.m. for 45 min at 4 °C. The fibril pellet was resuspended in PBS, pH 7.4. 10 μl of the fibril solution (10 μM) was spotted on carbon-coated Formvar EM grids and incubated at for 5 mins. The grid was washed gently with autoclaved milli-Q water. Freshly prepared and filtered 1% (v/v) uranyl formate solution (5 μl) was used for staining for 5 mins. The grids with samples were air-dried and imaged using a transmission electron microscope (Philips CM-200, Netherlands) with a magnification range of 6,600 X to 12,000X at 200 kV. Recording of images was done digitally with the aid of the Keen View Soft imaging system (Olympus, Japan). Further, for preparation of fibril seeds, the resuspended core fibrils were sonicated in a water bath for 3 mins at room temperature.

**Cell culture**

MCF 10A (non- tumorigenic human breast epithelial cells) used in the study was obtained from the cell repository at the National Centre for Cell Science, Pune, India. HFF cells (normal human foreskin fibroblasts) were obtained from ATCC. Both MCF 10A and HFF cells were used for experimentation at early passages. For culturing of MCF 10A, DMEM growth medium (Gibco, USA) supplemented with 5% horse serum (Gibco, USA), 0.5 μg/ml hydrocortisone (Himedia, India), 20 ng/ml hEGF (Invitrogen, USA), 10 μg/ml insulin (Himedia, India) and 1X Pen-Strep (Himedia, India) antimicrobial agent was used. Cells were not allowed to reach more than 80% confluency at each passage. For all experiments of MCF 10A, assay medium was used with the following composition: DMEM growth medium (Gibco, USA) supplemented with 2% horse serum (Gibco, USA), 0.5 μg/ml hydrocortisone (Himedia, India), 10 μg/ml insulin (Himedia, India) and 1X Pen-Strep antimicrobial agent (Himedia, India). For culturing HFF cells, DMEM growth medium supplemented with 5% fetal bovine serum (Gibco, USA), 10 mM HEPES buffer (Himedia, India) and 20 mg/ml gentamycin (Gibco, USA) as an antimicrobial agent was used.

**Fibril treatments of cells**

For core fibril treatment of MCF 10A, the assay medium was used for all the experiments. For that, 30 μM of Core fibril seeds were added to DMEM growth medium (Gibco, USA) supplemented with 2% horse serum (Gibco, USA), 0.5 μg/ml hydrocortisone (Himedia, India), 10 μg/ml insulin (Himedia, India) and 1X Pen-Strep antimicrobial agent (Himedia, India). For fibril treatment of HFF, 30 μM of Core fibril seeds were added to DMEM growth medium supplemented with 5% fetal bovine serum (Gibco, USA), 10 mM HEPES buffer (Himedia, India) and 20 mg/ml gentamycin (Gibco, USA) as antimicrobial agent. Cells were incubated for 48 hrs in a 5 % CO_2_ incubator at 37 °C.

**Immunocytochemistry**

Cells were treated with p53 core fibrils as discussed above and immunofluorescence was performed to study the state of p53 using p53 antibody. To do that, cells were fixed on the coverslips using 4% paraformaldehyde (Himedia, India) for 20 mins at room temperature. These fixed cells were washed with PBS, pH 7.4 and permeabilized using 0.2% Triton X (Sigma-Aldrich, USA) in PBS for 10 mins. The cells were then washed with PBS and incubated with 2% BSA (Himedia, India) in PBS for 1 hr to block non-specific epitopes. Cells were then incubated overnight at 4°C with mouse monoclonal anti-human p53 primary antibody DO-1 (Santa Cruz Biotechnology, Inc., USA) with 1:200 dilution. The cells were then washed thrice with PBST (phosphate buffer saline 0.1% Tween 20), pH 7.4. Then the coverslips were incubated with Goat anti-Mouse Alexa Fluor 488 conjugated secondary antibody (Invitrogen, USA) with 1:500 dilution for 2 hrs at room temperature in a moist and dark chamber. Cells were rinsed thrice with PBST, pH 7.4. The coverslips were mounted with mounting media containing 1% DABCO in 90% glycerol and 10% PBS. The slides were observed under the Zeiss Observer Z1 microscope (Zeiss, Germany) fitted with high-speed microlens-enhanced Nipkow spinning disc. Cells showing cytoplasmic and/or nuclear punctate were counted and the percentage of cells was plotted in each fraction. n=3 independent experiments (Total cells counted, N>200).

For the detection of Epithelial to Mesenchymal transition (EMT) markers, immunostaining of spheroids derived from MCF 10A cells (at T5 generation) was done. MCF 10A spheroids were grown on coverslips (details discussed in the relevant section below), fixed and permeabilized followed by blocking of non-specific epitopes as mentioned above. MCF 10A spheroids were incubated with 1:500 dilution of rabbit monoclonal EMT antibodies: Beta-catenin, Slug, Vimentin, N-cadherin (Santa Cruz Biotechnology, Inc., USA), overnight at 4ºC in a humidified chamber. Coverslips were washed three times with PBST, pH 7.4 and were further incubated with goat anti-rabbit Alexa 555 conjugated secondary antibody (1:1000) for 2 hrs at room temperature in a humidified chamber. MCF 10A spheroids were rinsed three times with PBST, pH 7.4. and stained with 1 μg/ml DAPI for 5 mins and were washed with PBS, twice. The coverslips were mounted and observed under Leica DMi8 (Leica, Germany) fluorescence microscope and images were analyzed using Image J software.

**Immunoprecipitation**

MCF 10A and HFF cells were grown and treated with core fibrils for 48 hrs. Immunoprecipitation was carried out as per the manufacturer’s protocol in non-denaturing conditions (Abcam, Cambridge, MA, USA). Briefly, cells were trypsinized and lysed under cold conditions using lysis buffer (20 mM Tris-HCl, pH 8.0, 137 mM NaCl, 1% NP-40, 2 mM EDTA) with protease inhibitor cocktail (Roche, Switzerland) under constant agitation for 20 min at 4 °C. The mixture was centrifuged at 10 000 r.p.m. for 20 min at 4 °C. The supernatant obtained was incubated with anti-p53 antibody (DO-1) under rotation at 4 °C for 8-10 hrs. Further, 120 μl of sepharose G beads were added to the solution and incubated at 4 °C under constant agitation for 4 hrs. This was centrifuged at 2500 r.p.m. for 30 secs to collect the beads. Elution was done with 100 μl of 0.2 M glycine buffer, pH 2.6 in 10 mins incubation with constant mixing and centrifugation at 2500 r.p.m. for 2 min at 4 °C. The eluted fractions were pooled and neutralized by adding an equal volume of Tris HCl, pH 8.0. The eluted protein was quantified using Bradford assay to ensure equal loading in the dot blot. For the immunoprecipitation of p53 from cell generations, cell pellets from sequential cell generations were collected for both fibril treated and untreated cells and immunoprecipitation of p53 was done as using the same method.

**Dot blot assay**

The immunoprecipitated p53 protein from MCF 10A and HFF cells was used for dot blot assay. For the detection of p53 amyloid aggregates in cells, 5 μg protein was used. For the passaging experiments, 3 μg of protein was used for the blots to prevent over-saturation of signals. To do dot blot, samples were spotted on nitrocellulose membrane and allowed to air dry. The membranes were then treated with a blocking solution of 5% non-fat skimmed milk powder (Himedia, India) in TBST for 1 h at 25 °C. These blots were washed with TBS and incubated with primary antibody (anti-p53, 1:200 dilution or OC, 1:500 dilution) overnight at 4°C on a rocker. After incubation, the blots were washed three times with TBST for 10 min each, followed by 1 h incubation with (anti-mouse for p53 and anti-rabbit for OC) horseradish peroxidase-conjugated secondary antibody, (ThermoFisher Scientific, USA) with 1:10000 dilution at room temperature on a rocker. Nonspecific binding was removed by three washes of TBST, each for 10 min. The signals were detected with Immobilon Western chemiluminescent HRP substrate (Merck Millipore, USA).

**Fluorescence-activated cell sorting (FACS) for cellular toxicity**

Both MCF 10A and HFF cells were seeded at a density of 10^4^ cells per well and incubated for 48 hrs in a 5 % CO_2_ incubator at 37°C. These cells were then treated with fibrils as mentioned above and further Actinomycin D (ActD, Sigma Aldrich, USA) was treatment was done. 2 μg/ml ActD was added for another 12 hrs to induce apoptotic stress. The cells were trypsinized and stained using an Annexin-V PI apoptosis kit according to the manufacturer’s protocol. For quantifying the apoptotic population, FACS analysis was done in BD FACS Aria SORP instrument (BD Biosciences, USA). n=3 independent experiments. The statistical significance was determined by one-way ANOVA followed by Bonferroni comparison post hoc test; p-values in the figure indicate *<0.05, ** <0.01, ***< 0.001.

**Soft agar colony formation assay**

Soft agar assay was done to study the transformation potential of core fibril treated MCF10A and HFF cells. Untreated cells were used as control. For soft agar assay, the 2X cell culture medium was prepared by dissolving 1 g of DMEM powder medium (GIBCO, USA) and 0.2 g of sodium bicarbonate (Himedia, India) in de-ionized water to a final volume of 50 ml. It was passed through a 0.2 μm filter to sterilize. 1.8% agarose was prepared by adding 1 g of noble agar to 100 ml of deionized water. Further, agarose was diluted to 0.3% and 0.6% using pre-warmed 2X culture DMEM medium for the bottom and top layers, respectively. The 12-well plates were covered with 0.3% agarose mixture (bottom layer), which was allowed to solidify at room temperature, in cell culture hood, for 30 mins. For the upper layer, fibril treated and untreated cells were harvested by trypsinization, centrifuged, resuspended in complete media. 0.6% agarose solution was melted in a microwave and mixed with cell suspension in a 1:1 ratio (at 42 °C), such that there are 10^6^ cells per well. This mixture was then layered on top of the bottom layer. The cell/agar mixture was allowed to solidify at room temperature, in cell culture hood, for 30 min before placing it into a 37°C humidified cell culture incubator. The plates were observed daily for the formation of colonies. The colonies were imaged under a bright-field microscope Leica DMi1 (Leica, Germany) and counted. n=3 independent experiments were done. The statistical significance was determined by one-way ANOVA followed by Bonferroni comparison post hoc test; p-values in the figure indicate *<0.05, ** <0.01, ***< 0.001.

**Detection of cell proliferation marker Ki-67**

MCF 10A and HFF cells (fibril treated for 48 hrs and untreated) were seeded at a cell density of 10^4^ cells per well on a glass coverslip. Cells were fixed, permeabilized and non-specific epitopes were blocked as mentioned above. Cells were then incubated overnight at 4°C with 1:1000 dilution of anti-Ki67 antibody-Alexa Fluor 488 tagged (Abcam, UK). Then cells were washed thrice with PBST (phosphate buffer saline 0.1% Tween 20), pH 7.4. The coverslips were mounted with mounting media containing 1% DABCO. The slides were observed under Zeiss Observer Z1 fitted with high-speed microlens-enhanced Nipkow spinning disc and the cells positive for the Ki-67 marker was counted for each set of samples (n>200). n=3 independent experiments. The statistical significance was determined by one-way ANOVA followed by Bonferroni comparison post hoc test; p-values in the figure indicate *<0.05, ** <0.01, ***< 0.001.

**Wound healing assay**

To determine the effect of p53 aggregation on cell motility (migration capability) wound-healing assay was performed (Rodriguez et al., 2005). To do that, MCF 10A or HFF cells were cultured on a 12-well plate and were treated with 30 μM core fibril seeds for 48 hrs as described before. Untreated cells were used as controls. Cells were then washed with PBS, pH 7.4 and scratched with a sterile plastic tip. The peeled off cells were removed with two PBS washes and fresh media was added to the cells. Mitomycin C (0.5 μg/ml) (Sigma-Aldrich, USA) was added as a cell proliferation inhibitor to calculate the migration rate accurately. Cells were further incubated for 48 hrs to allow wound coverage. Images at zero and final time-points were acquired under a bright-field microscope (Leica DMi1). Wound width was measured using the ImageJ software. Cell migration was calculated using the formula: rate of cell migration (μm/hr)= [(0) time wound width- final wound width)/ time taken (in hrs)]. n=3 independent experiments. The statistical significance was determined by one-way ANOVA followed by Bonferroni comparison post hoc test; p-values in the figure indicate *<0.05, ** <0.01, ***< 0.001.

**Cell survival assay across different generations**

MCF 10A and HFF cells were seeded in a 6 well plate at a cell density of 5 x 10^4^ cells per well. The cells were then treated with 30 μM p53 core fibril seeds for 48 hrs as described previously. Untreated cells were kept as control. The cells were allowed to reach confluency (~80%) before passaging them further. To do that ~1% cells from the previous generation mixed with fresh media were added to a fresh well to get the next generation. Untreated cells were passaged similarly. For estimating the cell viability at each generation, Annexin-V PI staining coupled with FACS analysis was done. For each sample, 10,000 cells were analyzed. The passaging of cells to obtain sequential generations was continued until the cells lost their adherence potential (along with low viability). During this passaging, at each generation, cells were also seeded on glass coverslips for immunostaining with p53 antibody as mentioned before to check the p53 status. n=2 independent experiments. The statistical significance was determined by one-way ANOVA followed by Bonferroni comparison post hoc test; p-values in the figure indicate *<0.05, ** <0.01, ***< 0.001. The cell pellet was collected at each generation for immunoprecipitation of p53 as discussed.

**Senescence assay**

MCF 10A cells were cultured in 24-well plates and treated with p53 core fibrils for 48 hrs and passaged, as discussed. The corresponding untreated cells were used as control. Senescence was evaluated at T1 and T5 generations. For this, the cell culture medium from 24-well plate was aspirated and the cells were washed with PBS (500 μl per well), twice. After washing, 250 μl of 4% PFA solution was used to fix the cells for 5 min at room temperature, followed by washing the cells two times for 5 min each with PBS. Then, to evaluate the senescent cell population, senescence-associated beta-galactosidase (SA-beta-gal) staining was done according to the established protocol (Gary and Kindell, 2005). Briefly, the SA-β-gal staining solution (pH 6.0) was prepared by mixing 0.1% X-gal (ThermoFisher Scientific, USA), 5 mM potassium ferrocyanide, 5 mM potassium ferricyanide, 150 mM sodium chloride and 2 mM magnesium chloride in 40 mM citric acid/sodium phosphate solution. 250 μl SA-β-gal staining solution was added to each of the wells containing fixed cells and the plate was incubated in the dark in a 37 °C incubator. After 72 hrs, the staining solution was removed and cells were washed with distilled water. The senescent cells contain galactosidase enzyme, which cleaves the chromogenic substrate X-Gal present in staining solution to develop a blue precipitate, which stains the senescent cells blue. Each well was observed under an inverted bright-field microscope Leica DMi1 (Leica, Germany) using a 10X objective and blue-stained cells were counted. The SA-β-gal positive cells are represented as a percentage of the total cell number (total cells, n>200) with three independent experiments. The statistical significance was determined by one-way ANOVA followed by Bonferroni comparison post hoc test; p-values in the figure indicate *<0.05, ** <0.01, ***< 0.001.

**Luciferase assay**

Luciferase assay was used for evaluating the p53 transcriptional activity in MCF10 A core fibril treated and untreated cells. For this assay, PG13 vector containing 13 repeats of p53 binding sites cloned upstream of firefly luciferase was used (Addgene # 16442) along with plasmid containing renilla luciferase (Addgene # 74444) as a control to normalize for translational variations across the samples. Cells were seeded (50,000 cells/well) in a 24-well plate and transfection was performed at 70 % confluency. To do that, 1 µg of plasmids (500 ng of PG-13 plasmid and 500 ng of renilla plasmid) were transfected per well using Lipofectamine 3000 (Invitrogen, USA). Cells were harvested after 48 hrs of transfection and luciferase levels were quantified using a dual luciferase assay according to manufacturer protocol (Promega, USA) in a Tecan microplate reader (Tecan, Switzerland). For rescue of p53 aggregation experiments, rescue peptide (10 µM) was added 16 hrs prior to harvesting the cells and the luciferase levels were measured as mentioned above. The ratio of firefly and renilla (relative luciferase activity) was used to indirectly evaluate the p53 transcriptional activity and the graphs were analyzed and plotted in GraphPad Prism 6.0. n=2 independent experiments. The statistical significance was determined by one-way ANOVA followed by Bonferroni comparison post hoc test; p-values in the figure indicate *<0.05, ** <0.01, ***< 0.001.

**MTT assay**

MTT assay was done to evaluate the cytotoxicity of the MCF 10A cells (fibril treated and untreated at T5 generation) in the presence and absence of various drugs. The drugs Cisplatin, Doxorubicin and Paclitaxel (Sigma-Aldrich, USA) were dissolved in DMSO to make the stock concentration of 1 mM and further dilutions of each drug were made in sterile PBS (pH =7.4). MCF 10A cells at T5 (untreated and core fibril treated) were independently seeded in a 96 well plate at a density of 10^4^ cells per well and incubated overnight. 100 μl of DMEM assay media containing the drugs of increasing concentration (0.5, 1, 2.5, 5, 10 and 20 µM for Cisplatin and Paclitaxel, 0.05, 0.10, 0.50, 1, 1.5, 2, 5,10, 20 µM for Doxorubicin) was added to the wells. The samples were done in triplicates. PBS buffer and DMSO was added to the media with cells as a buffer control (negative control). Triton X added to the media with cells was used as a positive control. The media with cells (100% viability) and cell-free media alone (for background absorbance) also were used. The plate containing cells and drugs were incubated for 24 hrs. After incubation, 10 μl of MTT dye (Sigma Aldrich, USA) (5 mg/ml) in PBS was added to the wells and incubated for 4 hrs. Then, 100 μl of DMF-SDS was added and incubated overnight. Further, the absorbance at 560 nm and background scattering at 690 nm was measured in a SpectraMax M2® (Molecular devices, USA) plate reader. The background scattering values at 690 nm were subtracted from the absorbance values at 560 nm to calculate the viability of cells both in the presence and absence of various compounds at each drug concentration. EC_50_ of each drug was calculated using GraphPad Prism 6. n=3 independent experiments. The statistical significance was determined by one-way ANOVA followed by Bonferroni comparison post hoc test; p-values in the figure indicate *<0.05, ** <0.01, ***< 0.001.

**Spheroid formation and calcein-AM staining**

MCF10A (core fibril treated and untreated cells) passaged cells at T5 generation were evaluated for their ability to form spheroids in Matrigel® (Corning Inc., USA). 10 µl of Matrigel® was used to layer the surface of clean glass coverslip placed in a 24 well plate and allowed to solidify. Fibril treated and untreated MCF 10A cells (10^3^ cells per well) were mixed individually with 50 µl Matrigel® and layered on the top of Matrigel bed on the coverslip. The Matrigel® was allowed to solidify before the addition of fresh media on the cells. The cells were incubated for 10 days and imaged periodically along with the untreated control to detect spheroid formation. The diameter of spheroids (D) was measured from the images of larger spheroids. Spheroid volume was calculated using the formula, the volume of spheroid= 4/3 X 3.14(r)^3^, where r is the radius and r=D/2. n=2 independent experiments. Following spheroid formation, the spheroids were stained with a mixture of two dyes: 1 μM calcein-AM (ThermoFisher Scientific, USA) and 1 μM of ethidium homodimer-1 (ThermoFisher Scientific, USA). Dyes were prepared freshly in sterile PBS and added directly to the media without aspiration and incubated for 2 hrs before imaging. The dye solution was washed out with PBS slowly to avoid spheroid loss, disintegration, or displacement. Spheroids were imaged using DMi8 microscope (Leica, Germany) with color HD camera

**Animal handling**

Female SCID mice (6-8 weeks old) weighing ~20–25 g were obtained from the ACTREC, Mumbai, India and acclimatized to the new location for a week. All animals were housed under standard conditions with a 12 hrs dark/light cycle, kept in micro-isolator cages with autoclaved bedding, with 50% humidity, a temperature of 22 ± 2°C and free access to water and food (standard pellet feed). The study (Approval certificate: ARI/IAEC/2019/03) was approved by the Institutional Animal Ethics Committee, Agarkar Research Institute, Pune.

**Mouse xenograft using fibril-transformed cells**

MCF 10A cells were treated with core fibrils and passaged till T5 generation as discussed. p53 aggregates in the passaged cells at T5 were confirmed by immunofluorescence. Untreated MCF 10A cells (non-tumorigenic) were parallelly passaged till T5 and did not show p53 aggregation. Both core fibril treated and untreated cells at T5 generation were used to analyze the tumorigenicity by injection in immunocompromised SCID mice. The mice were divided into groups of 4 for each experimental set. The groups were as follows: MCF 10A untreated cells, MCF10A core treated cells and MCF 7 cells. Each mouse received injection subcutaneously in the mammary fat pad with 2x10^7^ core transformed cells in sterile PBS (150 μl each injection). Untreated passaged cells (1x10^7^) were injected as a control. MCF 7 cells (1x10^7^ cells) were used as a positive control for tumor induction. Estradiol (Estrabet tablets, 2 mg, Abbott India Ltd, India) treatment was given orally to mice via their drinking water (1000 nM i.e. 0.054 µg/25 g mouse/day). All animals which received injections of MCF 10A core treated cells and injections of MCF 7 developed tumors. The mice were weighed once a week and the area of injection was observed. After tumor induction, the tumor volume was periodically measured using width (a) and length (b) measurements (a^2^b/2, where a < b), to assess the tumor growth. FDG coupled with positron-emission tomography (PET) and CT scan was done to study the tumor morphology and metabolism. At the end of the experiments, the tumors were excised and embedded in paraffin to be sectioned for characterization. The experiment was repeated twice with mice in groups of 4, each time.

**Micro-PET-CT acquisition**

The experimental mice injected with MCF10A (core fibril treated and untreated) were kept for overnight fasting with access to drinking water before^18^F FDG (2-deoxy-2-[fluorine-18] fluoro-D-glucose) injection. Animals were anesthetized by isoflurane and 25-30 MBq ^18^F-FDG (~800 microcuries) were injected intravenously through the tail vein. Whole-body positron emission tomography (PET) scan integrated with computed tomography (CT) scan was performed after ^18^F-FDG administration (within one hour of ^18^F-FDG administration). Both CT and PET scans were performed with the tri-modality gamma imaging system (Triumph@, Gamma Medica Ideas, Northridge, CA, USA). Image reconstruction has done by Maximum Likelihood Expectation Maximization (MLEM), an iterative algorithm. Reconstructed PET/CT images were analyzed using PMOD software, version 3.2 (PMOD Technologies Ltd, Zurich, Switzerland). ROIs were drawn for the tumor-bearing animals in the micro-PET image with the guide of the aligned micro-CT image. Image-based method was used to calculate the standard uptake values (SUV) of ^18^F-FDG in the tumors considering the body weight and injected a dose of ^18^F-FDG. The SUV was calculated using the following formula: SUV= (Activity in Region of Interest)/ (Injected Dose/Body Weight).

**Hematoxylin and eosin (H & E) staining**

To compare the histology of tumor tissue (MCF10A core fibril treated and MCF 7) and normal tissue (only MCF 10A and RP treated core transformed MCF10A), hematoxylin and eosin staining was performed. Tissues were formaldehyde fixed and paraffin-embedded. Thin sections of the paraffin-embedded tissues were prepared using the microtome. The tissues were then deparaffinized using decreasing concentrations of the clearing agent xylene (100% xylene, followed by 50% xylene in 50% ethanol) and further rehydrated in decreasing concentrations of graded ethanol (100%, 95%, 70% and 50%) and finally with distilled water. The sections were then washed thoroughly with distilled water for 10 mins and then stained with 0.5% hematoxylin prepared in distilled water for 2 mins. The unbound stain was removed using 70% and 90% ethanol wash for 10 mins each. Next, 0.8% eosin in 95% ethanol was used to stain the sections for 1 min and kept in xylene for 1 hr. The sections were then sealed with a coverslip using DPX mounting solution (a mixture of polystyrene, tricresyl phosphate and xylene) and observed under a DMi8 microscope (Leica, Singapore) with color HD camera.

**Immunohistochemistry of tissues**

Paraffin-embedded tissues, tumor tissue (MCF 10A core fibril treated and MCF 7) or normal tissue (only MCF 10A and RP treated core transformed MCF 10A) were used for immunohistochemistry. Thin sections of the paraffin-embedded tissues were prepared using a microtome. The tissues were then deparaffinized using decreasing concentrations of the clearing agent xylene (100% xylene, followed by 50% xylene in 50% ethanol) and further rehydrated in decreasing concentrations of graded ethanol (100%, 95%, 70% and 50%) and finally with distilled water. Enzymatic antigen retrieval was performed on the sections by treating them with 0.05% trypsin at 37ºC for 2 mins. Sections were then washed with TBST (Tris-buffered saline with tween-20) at pH 7.4 and treated with 0.2% Triton-X 100 in TBST for 10 mins for permeabilization of tissues. Nonspecific antigenic sites were blocked using TBST containing 2% BSA. First, the sections were incubated with mouse p53 antibody (DO-1) (1:200 dilution) and OC (1:500 dilution), overnight at 4ºC in a humidified chamber. The tissue sections were then washed three times with TBST, pH 7.4. The sections were then incubated with goat anti-mouse Alexa 488 conjugated secondary antibody (1:1000) and goat anti-rabbit Alexa 555 conjugated secondary antibody (1:1000) for 2 hrs at room temperature in a humidified chamber. Sections were then rinsed three times with TBST, pH 7.4. The sections were then stained with 1 μg/ml DAPI for 3 mins and were washed with TBS, twice. The sections were mounted with mounting media containing 1% DABCO (Sigma-Aldrich, USA) in 90% glycerol and 10% phosphate-buffered saline (PBS). Control tissue sections were stained using the procedure mentioned above. Sections were observed under Leica Mi8 fluorescence microscope (Leica, Germany) and images were analyzed using Image J software.

**Real-time PCR**

MCF 10A cells were treated with core fibril seeds and cultured till T5 generation as discussed. At each generation (T1 to T5), the cell pellet was collected for RNA isolation. RNA isolation was done from the treated cells and untreated cells using TriZol (Invitrogen, USA) method according to the manufacturer’s protocol. Briefly, the cell pellet was lysed using 700 μl of TriZol for a single 12-well. Further, 400 μl of chloroform was added and samples were vortexed vigorously. Samples were centrifuged at 12,000 g for 15 mins at 4°C. The aqueous phase (upper layer) was transferred to fresh tubes and an equal volume of isopropyl alcohol was added for precipitation of RNA. Samples were incubated at room temperature for 20 mins and centrifuged at 12,000 g for 10 mins at 4°C. The supernatant was discarded and RNA pellet was washed using 70% ethanol followed by centrifugation at 7,500 g for 5 mins at 4°C. Ethanol was allowed to evaporate and RNA was resuspended in nuclease-free water. The concentration of the isolated RNA was measured in a nanodrop spectrophotometer (Implen, USA). cDNA synthesis was done with RevertAid first-strand cDNA synthesis kit (ThermoFisher Scientific, USA) according to the manufacturer’s protocol with random hexamer primers. Real-time PCR (qRT-PCR) was performed using the SYBR Green method using primers for p21, BAX, DDB2, CCND2, MAPK1 genes (Refer Supplementary Table 1 for primer sequences). Maxima SYBR Green/ROX qPCR Master Mix (2X) (ThermoFisher Scientific, USA) was used according to the manufacturer’s protocol. n=2 independent experiments.

**Microarray analysis**

MCF 10A cells were treated with core fibril seeds along with untreated control and were cultured till T5 generation as described before. RNA isolation was done for T1 and T5 generation for fibril treated and untreated cells using the RNeasy mini kit (Qiagen, Germany) as per the manufacturer’s protocol. The concentration and purity of the RNA were evaluated using the Nanodrop Spectrophotometer (Thermo Scientific; 2000). The integrity of the RNA was analyzed on the Bioanalyzer (Agilent; 2100 expert). Microarray analysis (outsourced) was done by using RNA isolated from the first and last passage after treatment. The microarray hybridization and scanning were performed at the Agilent certified microarray facility of Genotypic Technology, Bengaluru, India. The samples for Gene expression were labeled using Agilent Quick-Amp labeling Kit (p/n5190-0442). The total RNA was reverse transcribed at 40°C using oligo dT primer tagged to a T7 polymerase promoter and converted to double-stranded cDNA. Synthesized double-stranded cDNA was used as a template for cRNA generation. cRNA was generated by *in vitro* transcription and the dye Cy3 CTP(Agilent) was incorporated during this step. The cDNA synthesis and *in vitro* transcription steps were carried out at 40°C. Labeled cRNA was cleaned up using Qiagen RNeasy columns (Qiagen, Cat No: 74106) and quality assessed for yields and specific activity using the Nanodrop ND-1000. Labeled cRNA samples were fragmented at 60ºC and hybridized on to an Agilent Human Gene Expression Microarray 8X60K. Fragmentation of labeled cRNA and hybridization were done using the Gene Expression Hybridization kit of (Agilent Technologies, *In situ* Hybridization kit, Part Number 5190-0404). Hybridization was carried out in Agilent’s Surehyb Chambers at 65º C for 16 hrs. The hybridized slides were washed using Agilent Gene Expression wash buffers (Agilent Technologies, Part Number 5188-5327) and scanned using the Agilent Microarray Scanner (Agilent Technologies, Part Number G2600D). Raw data extraction from images was obtained using Agilent Feature Extraction software. Feature extracted raw data was analyzed using Agilent GeneSpring GX (v14.5) software. Normalization of the data was done in GeneSpring GX using the 75th percentile shift method [Percentile shift normalization is a global normalization, where the locations of all the spot intensities in an array are adjusted]. This normalization takes each column in an experiment independently, and computes the percentile of the expression values for this array, across all spots (where n has a range from 0-100 and n=75 is the median). It subtracts this value from the expression value of each entity and fold change values were obtained by comparing test samples with respect to specific control samples. Significant genes upregulated with fold change >=1 (logbase2) and down-regulated with fold change

<=-1 (logbase2) in the test samples with respect to the control sample were identified. p-values were calculated using Benjamini-Hochberg correction for multiple hypothesis testing to calculate the false discovery rate (FDR). Genes with more than 2-fold change in gene expression and an adjusted p-value (False discovery rate (FDR)) <0.05 were classified as differentially expressed and used for further analysis. Differentially regulated genes were clustered using hierarchical clustering based on the Pearson coefficient correlation algorithm to identify significant gene expression patterns. Biological analysis was performed for the differentially expressed genes based on their functional category and pathways using Metascape (Zhou et al., 2019b) (http://metascape.org/).

**Sample preparation for proteomics analysis**

Proteomic analysis was carried out using iTRAQ labeling coupled with LC-MS/MS (Datta et al., 2017; Zhao et al., 2016). For this, MCF 10A cells were treated with core fibril seeds and cultured till T5 generation as described before**.** The cells without core fibril treatment were used as control. Cells at T1 and T5 passages were pelleted down at 4°C. Proteins were extracted from the harvested cell pellet by RIPA lysis and extraction buffer (Sigma-Aldrich; 150 mM NaCl, 1.0% IGEPAL^®^ CA-630, 0.5% Sodium deoxycholate, 0.1% SDS, 50mM Tris, pH 8.0) containing protease and phosphatase inhibitor cocktail (ThermoFisher Scientific, USA) with intermediate vortexing, followed by mild sonication. The lysate was centrifuged at 14,000 × g for 30 mins at 4°C. The supernatant was collected and acetone precipitation of the proteins was performed at −20°C. The protein precipitates were re-solubilized in 8 M urea. The protein concentration was estimated by BCA protein assay kit (ThermoFisher Scientific, USA).

**iTRAQ labeling and fractionation by reverse phase chromatography**

From each sample, an equal amount of protein (40 μg) was reduced with 5 mM DTT (37°C, 60 min) and alkylated with 20 mM iodoactamide (IAA) (room temperature, 30 min, dark). MS grade trypsin (Pierce, Thermo Fisher Scientific, USA) was added to convert the proteins to peptides (trypsin: protein ratio of 1:20 (w/w) at pH 8, 37°C for 24 h). The pH of the solution was lowered with 1% formic acid to quench the activity of trypsin. Further, the peptides obtained were labeled with isobaric mass tags (iTRAQ labeling) using iTRAQ Reagent 8-Plex kit (SCIEX, USA) based on the manufacturer's protocol. In the T1 generation, the samples were labeled with isobaric mass tags as follows: control (untreated) samples with mass tags 113 and 115, P8 peptide treated (Ghosh et al., 2017) samples with mass tag 114 (data not shown) and core fibril treated samples with mass tag 116. Similarly, in the T5 generation the samples were labeled as follows: control (untreated) samples with mass tags 117 and 119, P8 peptide treated samples (data not shown) with mass tag 118 and core fibril treated samples with mass tag 121. The samples were incubated at room temperature for 2 hrs before pooling them together. Two biological repeats were carried out. To reduce the proteome complexity, basic reverse phase chromatography (bRP) was performed using 1260 Infinity HPLC system (Agilent, USA). A total of 320 μg combined labeled peptides were loaded onto the C18 column (Agilent, USA, 300 extend-C18; 3.5µm; 2.1 × 150 mm) which was previously equilibrated with solvent A (10 mM ammonium formate, pH 10) and the column temperature was maintained at 40°C. The peptides were eluted with the gradient from 2-50% of solvent B (10 mM ammonium formate in 90% acetonitrile, pH 10) over 65 min total run at a flow rate of 0.5ml/min. The eluted fractions were pooled to 10 fractions for each biological replicate, vacuum dried, desalted with C18 tips (Pierce, USA), and reconstituted with solvent C (2% (v/v) acetonitrile, 0.1% (v/v) formic acid in water) for LC-MS/MS analysis.

**LC-MS/MS acquisition**

All HPLC purified fractions were analyzed in Sciex 5600^+^ Triple-TOF mass spectrometer coupled with ChromXP reversed-phase 3 μm C18-CL trap column (350μm × 0.5 mm, 120 Å, Eksigent) and nanoViper C18 separation column (75 μm × 250mm, 3 μm, 100 Å; Acclaim Pep Map, Thermo Scientific) in Eksigent nanoLC (Ultra 2D plus) system. The binary mobile solvent system was used as follows: solvent C (2% (v/v) acetonitrile, 0.1% (v/v) formic acid in water) and solvent D (98% (v/v) acetonitrile, 0.1% (v/v) formic acid). The peptides were separated using a 70 min gradient from 5-50% of solvent D at a flow rate of 200 nl/min. The MS data of each fraction was acquired in IDA (information-dependent acquisition) with high sensitivity mode. The collision energy was set to iTRAQ reagent. Each cycle consisted of 250 and 100 ms acquisition time for MS1 (m/z 400−1250 Da) and MS/MS (70–1800 m/z) scans respectively with a total cycle time of 2.3 s. Each fraction was run in duplicate.

**Peptide and Protein Identification**

All raw files (.wiff) were converted to Mascot generic file (mgf) format using Peak View (version 1.2.0.3). ProteinPilot software (version 4.5, SCIEX) with the Paragon algorithm was used for protein identification and relative iTRAQ quantification. Proteins were identified against the UniProt human-reviewed database containing only canonical sequences (downloaded in April 2020). The search parameters were set as follows: iTRAQ 8plex (peptide labeled); IAA Cysteine alkylation; digestion enzyme Trypsin. Peptides and proteins were validated at ˂1% false discovery rate (FDR) and with unused score ˃ 1.3 (which corresponds to ˃ 95% confidence). The cutoff value for up-regulation and down-regulation of proteins was set to ˃ 1.3 and ˂ 0.8 respectively with p-value ˂ 0.05. The fold change was presented as the relative expression ratio of a given protein in the core fibril seeds treated cells to the untreated control. The differential abundance of proteins from two biological replicates were combined and used further for downstream analysis. The mass spectrometry proteomics data has been deposited to the ProteomeXchange Consortium via PRIDE partner repository with the dataset identifier PXD019498.

**Gene Ontology Analysis (Pathways and Functional Enrichment)**

The list of differentially expressed (the ratio threshold*>*1.3 or <0.80, p-value *<*0.05) proteins were extracted with Swiss-Prot accession. The web server-based software WEB-based Gene SeT AnaLysis Toolkit (Zhang et al., 2005) (http://webgestalt.org) and NetworkAnalyst 3.0 (Zhou et al., 2019a) were used with their default settings for functional enrichment and annotation. An overrepresentation enrichment analysis (ORA) was performed in WebGestalt for gene ontology (GO) analysis using a non-redundant functional database. Reactome database was used for pathway analysis. Signaling networks were built by using NetworkAnalyst 3.0 using the STRING interactome database (Szklarczyk et al., 2018).

**Rescue peptide (RP) treatment**

A cell-penetrating peptide, called rescue peptide (RP), previously established by Soragni et al, was used to disaggregate p53 amyloids (Soragni et al., 2016). The RP was custom synthesized by USV Limited, Mumbai, India by solid-state peptide synthesis method. Lyophilized RP was dissolved in DMSO to make a stock concentration of 10 mM. MCF 10A cells were treated with core fibrils in a 12 well plate, as discussed, to induce p53 aggregation. These cells were passaged till T5 generation. Untreated control was passaged parallelly as a control. 10 μM working concentration of the RP was used to treat MCF 10A core fibril transformed cells for 12 hrs. Fibril transformed cells (at T5) without RP treatment and untreated MCF 10A cells (at T5) were kept as control. The rescue peptide treated cells were used for soft agar (n=3 independent experiments), apoptosis (n=3 independent experiments) and p53 luciferase reporter assays for understanding the effect of p53 disaggregation (n=2 independent experiments) as described previously. To check the tumorigenic potential of RP treated MCF 10A core fibril transformed cells (at T5), the mice were divided into groups of 3 for each experimental set. The mice were injected subcutaneously in the mammary fat pad with 2x10^7^ cells (fibril transformed cells (at T5) with/without RP treatment and untreated MCF 10A cells (at T5)) in sterile PBS (150 μl each injection). The mice were maintained and tumor induction was monitored as described previously. FDG coupled with positron-emission tomography (PET) and CT scan was done similar to the previous experiment. Tissue sectioning along with H&E staining for tumor and normal tissue was done as described previously.

**Statistical Analysis**

The statistical significance (p-value) was calculated by one-way ANOVA followed by Bonferroni Multiple Comparison post hoc test. The p-value for the significance is *p ≤ 0.05, **p ≤ 0.01, ***p ≤ 0.001; non-significant (NS p > 0.05). KaleidaGraph, version 4.1 software was used for calculating the statistical significance.

**Supplementary Figures**

**Supplementary figure 1**

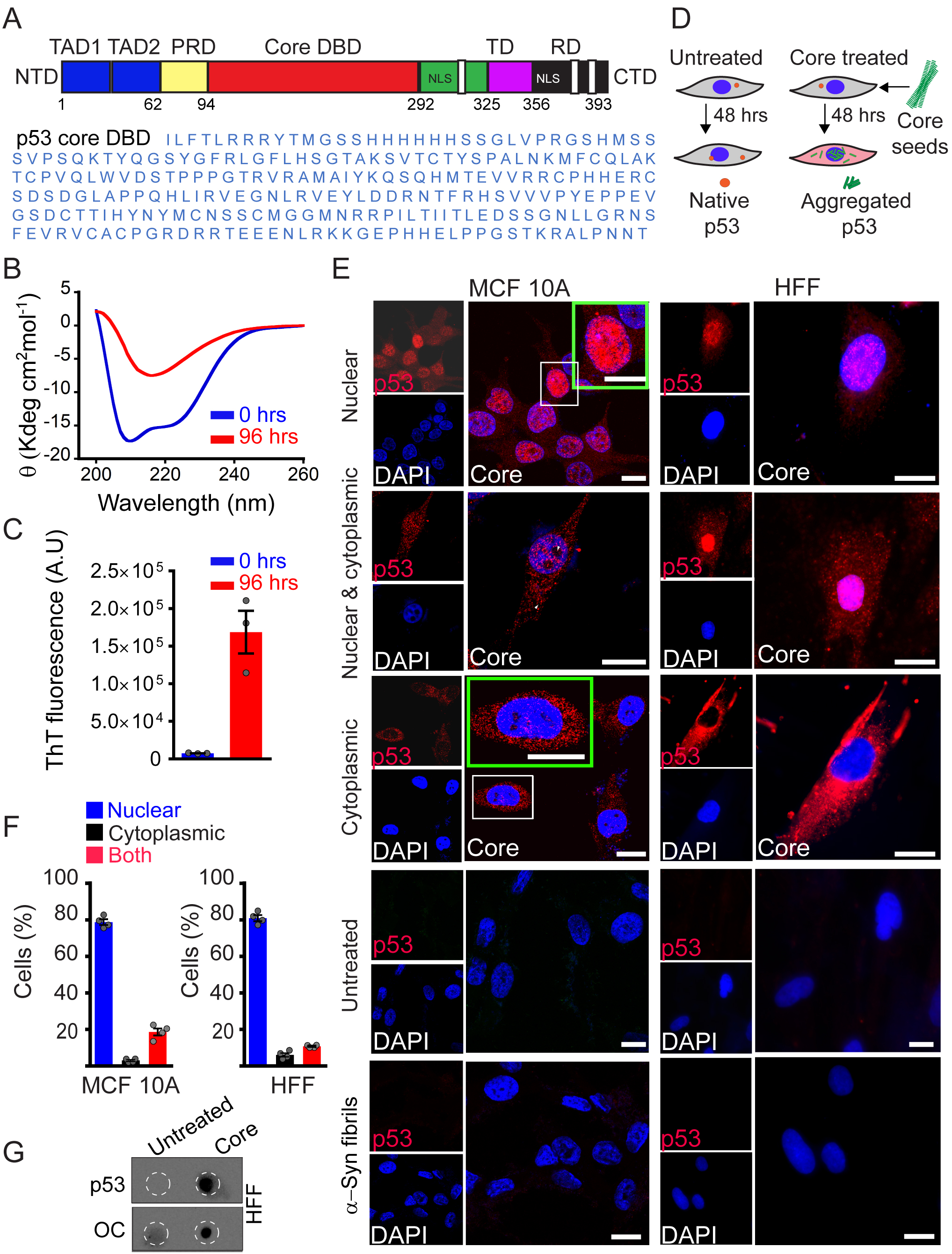

**Figure S1. *In vitro* amyloid formation of p53 core domain and induction of p53 amyloid formation in cells.** (A) Domain organization of full-length p53 tumor suppressor protein with 393 amino acid residues. NTD: N-terminal domains, TAD1 and TAD2: Transactivation domains 1 and 2, PRD: Proline-rich domains, Core DBD: core DNA-binding domain, TD: Tetramerization domain, RD: Regulatory domain, CTD C-terminal domains, NLS: Nuclear localizing sequence (upper panel). The amino acid sequence of the p53 core domain (lower panel). (B, C) Amyloid formation by p53 core domain. p53 core domain (50 μM) was incubated in the presence of an equimolar concentration of chondroitin sulfate (CSA), pH 7.4 at 37°C with agitation. The amyloid formation was measured using ThT fluorescence and CD spectroscopy after 96 hrs of incubation. (B) CD spectroscopy of p53 core domain protein at 0 hrs showing the characteristic α-helical structure, which showed the conformational transition to β-sheet after 96 hrs of aggregation. n=3 independent experiments. (C) ThT fluorescence at 480 nm showing higher ThT binding for p53 core after 96 hrs of incubation in the presence (CSA), indicating aggregation of p53 core domain. The values in the plots represent mean ± s.e.m, n=3 independent experiments. (D) Schematic representation showing fibril treatment of cells for 48 hrs, which leads to aggregation of intracellular p53. Sonicated p53 core fibrils (seeds) were added to MCF 10A and HFF cells followed by 48 hrs to induce p53 amyloid formation in the cells. (E) Immunostaining of p53 fibril seed treated cells showing p53 staining as punctate upon fibril treatment. Exogenous addition of p53 core fibrils induces p53 amyloid formation in cells. MCF 10A and HFF cells were treated with p53 fibril seeds for 48 hrs and immunofluorescence study was done using p53 antibody (DO-1). The localization of p53 aggregates observed in the nucleus as well as the cytoplasm (both the cells are represented). Immunofluorescence of cells without any fibrils treatment (untreated) or with treatment with α-synuclein fibril seeds for 48 hrs showing no significant p53 stabilization. This indicated that endogenous p53 aggregation is specific to the addition of p53 core fibril seeds. Scale bars are 10 μm. (F) Quantification of fibril treated cells showing the percentage of cells with nuclear and/or cytoplasmic p53 aggregates. The calculation was done using more than 200 cells (n> 200). For MCF 10A cells, ~79% of the population of cells showed only nuclear aggregates, ~3% of cells showed only cytoplasmic aggregates and ~18% of the cells showed both nuclear and cytoplasmic aggregates. For HFF cells, ~83% of the cell population showed only nuclear aggregates, ~7% of cells showed only cytoplasmic aggregates and ~10% of the cells showed both nuclear and cytoplasmic aggregates. The values plotted represent mean ± s.e.m, n=4 independent experiments. (G) Immunoprecipitation of p53 was done from the core fibril treated HFF cells. Dot blot analysis using OC amyloid fibril specific antibody confirmed the presence of p53 amyloids in the core fibril treated cells whereas untreated cells showed no p53 amyloid. n=2 independent experiments.

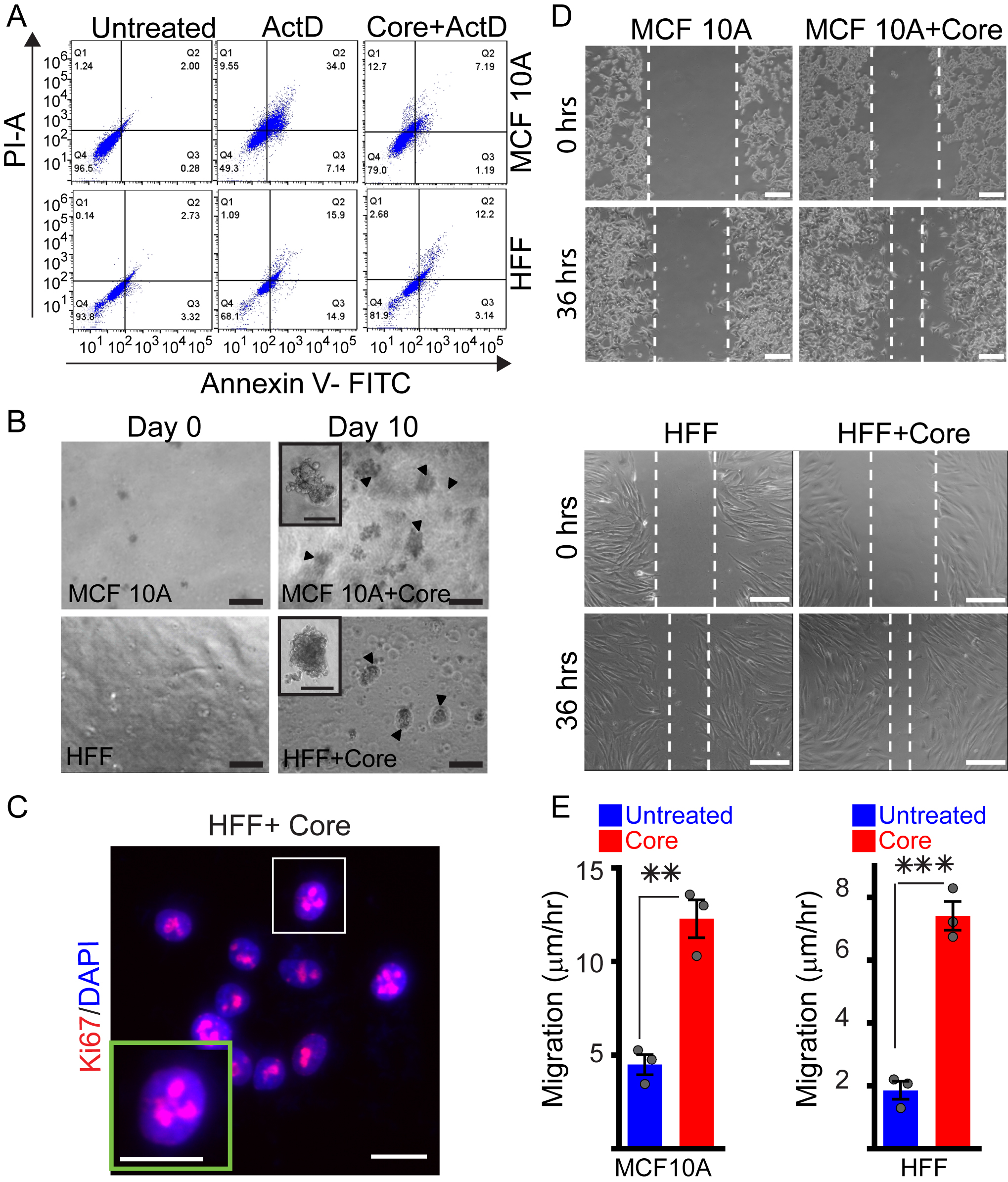
**Supplementary figure 2**

**Figure S2. p53 amyloid formation leading to loss of tumor-suppressive function and gain of oncogenic functions.** (A) Effect of p53 amyloid formation in drug-induced apoptosis. MCF 10A and HFF cells were pre-treated for 48 hrs with core fibril seeds followed by ActD. Untreated cells (control) and cells treated only with ActD (+ ActD) were used as control. Quantitative cell death assay using Annexin V PI coupled with flow cytometric analysis showing a high level of apoptosis in cells treated only with ActD (+ActD). However, core fibrils treatment before ActD treatment (Core+ActD) showed a marked decrease in apoptotic populations for both the cells. (B) Gain of colony formation ability by fibril treated cells. Soft agar colony formation assay showing the increased colony formation by both MCF 10A and HFF cells with core fibril treatment in contrast to untreated cells. Scale bars are 100 µm. (Inset scale: 50 µm). n=3 independent experiments. (C) Effect of p53 amyloid formation on the proliferation of HFF cells. Immunostaining of Ki67 (proliferation marker) showing fibril treated HFF cells with nuclear localization of Ki67. n=3 independent experiments. (D, E) Wound healing assay to examine the cell motility of fibril treated cells over 36 hours. (D) Microscopic image of scratch assay showing enhanced cell migration by MCF 10A and HFF when treated with core fibrils compared to untreated cells. Scale bars are 150 µm. (E) Quantification of cell migration of core fibril treated MCF 10A and HFF cells compared to the corresponding untreated control cells. Significantly enhanced cell migration was seen in cells treated with core fibrils seed as compared to untreated cells, for both MCF 10A and HFF cells. The values plotted represent mean ± s.e.m, n=3 independent experiments. The p values for MCF 10A and HFF core treated cells with respect to untreated cells are 0.00244 and 0.0005, respectively. Statistical significance (***p ≤ 0.001, **p ≤ 0.01) is determined by one-way ANOVA followed by Bonferroni multiple comparison post hoc test with 95% confidence interval.

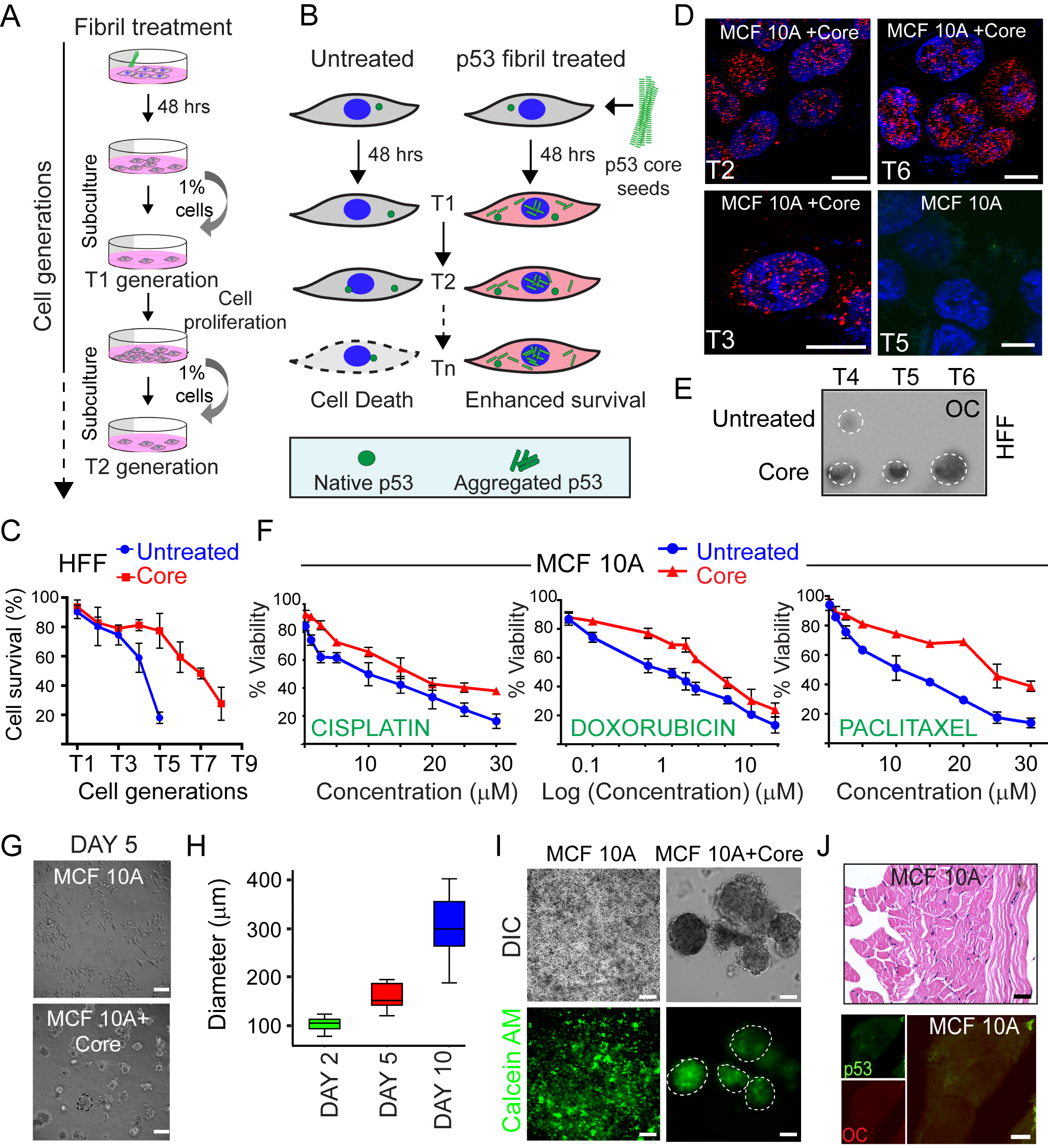
**Supplementary figure 3**

**Figure S3. Gain-of-function properties of fibril treated cells.** (A) Schematic representation of the survival experiment for cells treated with core fibrils. Cells were treated with p53 core fibrils seeds for 48 hrs and were allowed to reach confluency (~80%) before passaging them further. ~1% of cells from the previous generation were mixed with fresh media and added to a fresh well to get the next generation. Untreated cells were passaged similarly. Each cell generation after fibril treatment was labeled as T1, T2 and so on. (B) Schematic showing enhanced survival of the cells due to p53 core fibrils treatment. Cells after fibril treatment were used to obtain sequential cell generations. Untreated cells were correspondingly passaged. Cells with p53 aggregation showed enhanced survival of the cells as compared to untreated cells. (C) Cell viability of HFF fibril treated and untreated cells was calculated for all the generations (passages) using Annexin-V PI assay coupled with FACS analysis. The fibril treated HFF cells showing enhanced survival as compared to untreated cells. The values in the plots represent mean ± s.d, n=2 independent experiments. (D) Immunofluorescence staining of MCF 10A using p53 DO-1 antibody showing the presence of p53 aggregates through different passages (T2, T3 and T6 are shown) whereas the untreated control (T5 passage) did not show any p53 stabilization. Scale bars are 10 µm. (E) Immunoprecipitation of p53 from the fibril treated and untreated HFF cells followed by dot blot analysis using OC (amyloid fibril specific) antibodies. The OC positive immunoreactivity suggesting the presence of p53 amyloid aggregates in the fibril treated HFF cells at different passages. Untreated cells did not show the presence of p53 amyloid aggregates. n=2 independent experiments. (F) Drug resistance in core fibril treated cells. Dose-dependent toxicity of cells using MTT assay in the presence or absence of various drugs showing drug resistance in fibril treated cells compared to untreated cells. Increasing concentrations of cisplatin, doxorubicin and paclitaxel were used and cell viability was plotted against various concentrations of drugs. For doxorubicin, nanomolar concentration of drug was required for cytotoxicity, therefore, cell viability was plotted against log (concentration of doxorubicin) for better visualization. The values plotted represent mean ± s.e.m, n=3 independent experiments. (G-I) Spheroid formation by core fibril treated MCF 10A cells. (G) Three-dimensional cell culture using Matrigel by MCF 10A cells treated with or without p53 core fibril seeds. Day 5 data showing initiation of spheroid formation by core fibril treated cells; whereas no spheroid formation was observed for untreated cells. Scale bars are 150 µm. (H) The diameter of spheroids formed by core fibril treated MCF 10A cells, monitored over 10 days, showing a gradual increase of spheroid-size over time. Untreated cells did not show spheroid formation. The values plotted were representing the mean ± s.e.m, n=3 independent experiments. (I) The cell viability of spheroid formed by core fibril treated MCF 10A cells using Calcein-AM staining showing the viability of cells inside the spheroids. Scale bars are 150 µm. (J) Characterization of animal tissue sections using hematoxylin and eosin (H & E) staining and immunohistochemistry with p53 antibody. H & E staining of control tissue section where only MCF 10A cells (T5) were injected showing normal tissue morphology with a normal nucleus to cytoplasm ratio (upper panel). Scale bar is 100 µm. Immunohistochemistry of animal tissue using p53 antibody (DO-1) and amyloid antibody (OC) showing no p53 stabilization and amyloid formation, when dissected from the site where MCF 10A cells (untreated, T5) were injected (lower panel). Scale bar is 100 µm.

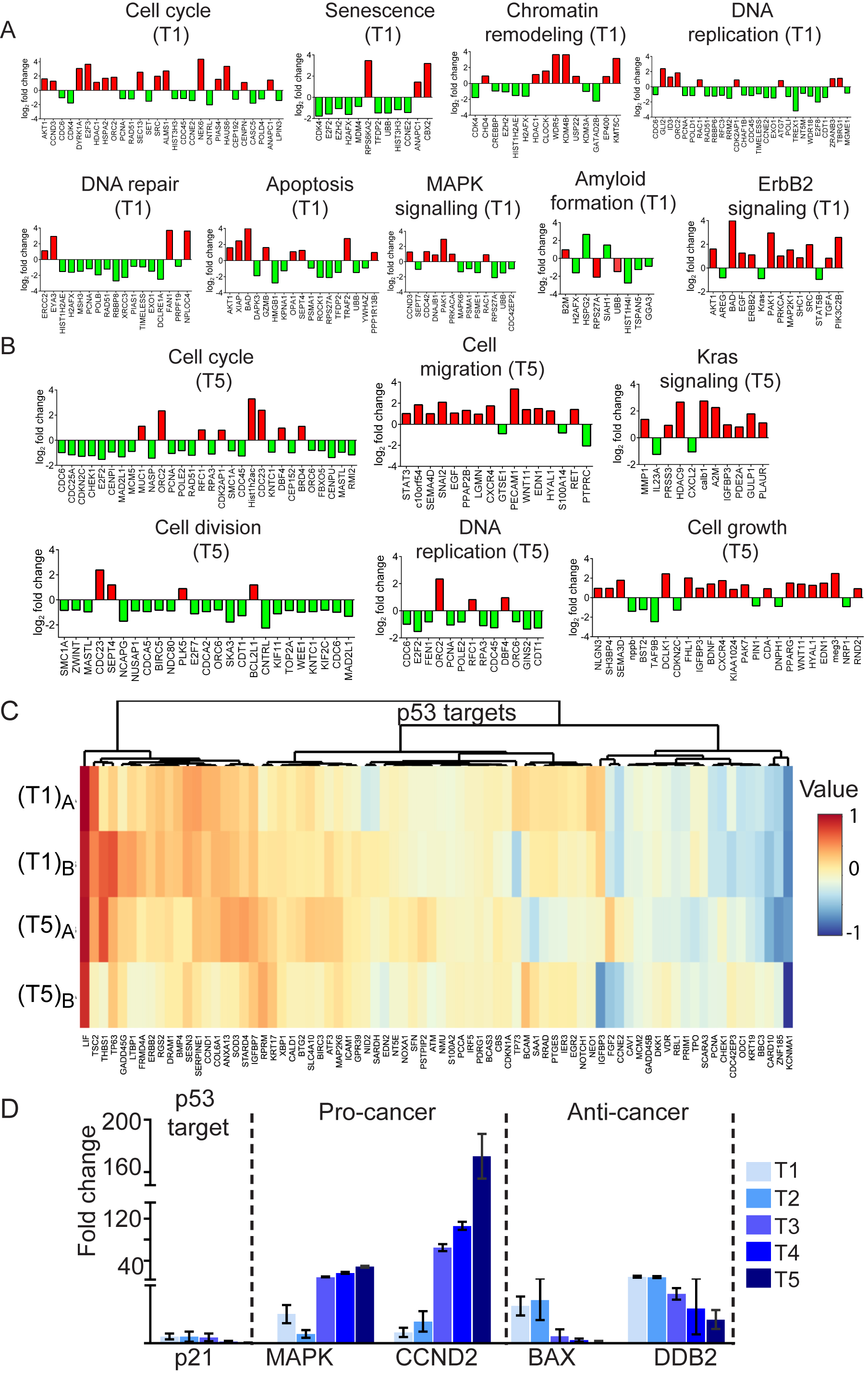
**Supplementary figure 4**

**Figure S4. Pathway analysis for gene expressions affected by p53 amyloid formation using microarray.** Gene ontology analysis showing differentially affected pathways due to p53 amyloid formation. Log_2_ fold change of some significantly altered genes from each pathway is plotted. The upregulated genes are shown in red bars whereas the down-regulated genes are shown in green bars. (A) Fold change of genes at T1 passages showing alteration in gene expression associated with various cellular processes such as cell cycle, apoptosis, DNA repair, DNA replication, senescence, chromatin remodeling, ErbB2 and MAPK signaling. This indicated that p53 amyloid formation at an earlier stage affects the tumor-suppressive functions of p53 by the cells. (B) Fold change of genes at T5 passages showing alteration of gene expression associated with cell cycle, cell growth, cell division, cell migration, Kras signaling and DNA replication. This deregulation of pathways might enhance the oncogenic phenotype of the cells by upregulation of oncogenes and suppression of anticancer genes. (C) Alteration of gene expression of p53 targets due to p53 amyloid formation. Microarray data showing differential gene expression of p53 targets in terms of heat map for core fibrils treated cells as compared to untreated cells. Specific gene targets of p53, which are affected upon p53 amyloid formation are shown. Two biological replicates are shown for each passage: (T1)_A_ and (T1)_B_ for T1 generation and (T5)_A_ and (T5)_B_ are for T5 generation. (D) Quantitative RT-PCR data showing the extent of specific gene expression during various generations. Expression of genes linked with p53 pathways such as p21, pro-cancer genes (MAPK and CCND2) and anti-cancer genes (BAX, DDB2) are analyzed. Expression of p21 decreased over the passages whereas pro-cancer genes were upregulated. The anti-cancer genes showed decreased expression over the passages indicating that p53 amyloid formation modulates the expression of genes, thereby enhancing the tumorigenic potential of the cells. The values plotted represent mean ± s.d, n=2 independent experiments.

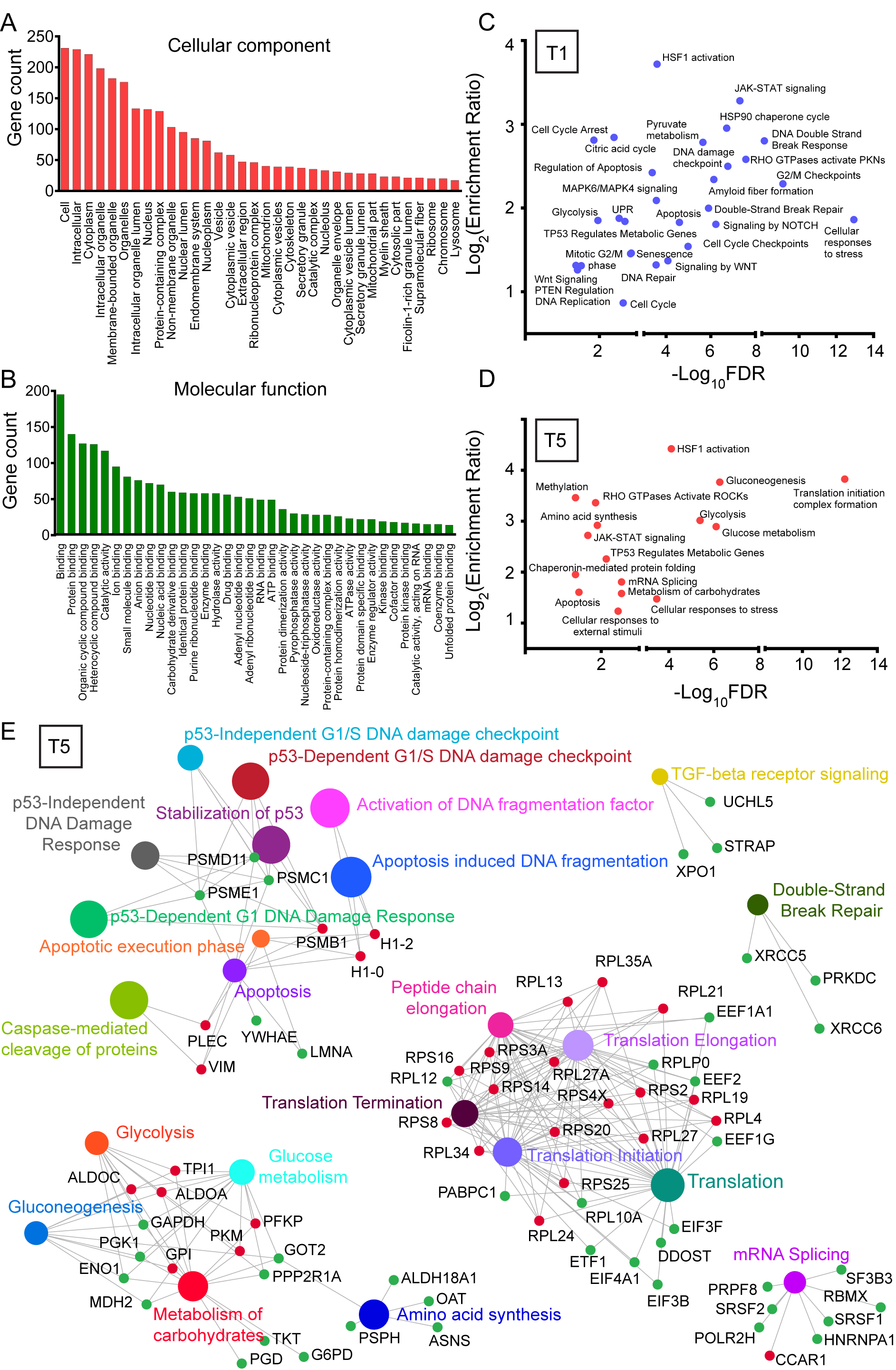
**Supplementary figure 5**

**Figure S5. Proteomic profiling of cells with p53 aggregates** **at T1 and T5 passages.** (A) and (B) Enrichment analysis of significantly dysregulated proteins in response to p53 amyloid formation at both T1 and T5 generations. Significantly affected proteins were annotated in terms of the cellular components (A) and the molecular functions (B). Most represented categories in the case of cellular components were intracellular category whereas most represented molecular functions include protein binding. (C) and (D) False discovery rate (FDR) and enrichment ratio of differentially expressed proteins with significantly altered signaling pathways were calculated from “Reactome database”. Log_2_(enrichment ratio) was plotted against the -log_10_(FDR) was plotted for both T1 and T5 passages. (C) Proteins enriched at T1 passage show that apoptosis, cell cycle, senescence and DNA damage repair pathways are significantly affected. Furthermore, the pathways linked to glucose metabolism are substantially upregulated, possibly contributing to higher survival of the cells. (D) At T5, cells harboring p53 aggregates showing significant changes in apoptotic pathways. Several signaling pathways (like JAK-STAT and Ras) that are altered upon p53 amyloid formation are also shown. The pathways involved in cellular metabolism like amino acid synthesis and carbohydrate metabolism showing upregulation, suggesting the highly metabolic state of these cells. (E) Analysis of significantly altered protein networks using NetworkAnalyst 3.0 showing dysregulated pathways upon fibril treatment to the cells. In the network, proteins are represented as nodes and clusters of proteins are represented as dense circles. At T5, p53 governed processes like apoptosis and DNA damage checkpoints show significant changes. The proteins associated with metabolic pathways and translational machinery also show increased expression, reminiscent of cancer cells. The upregulation of gain of function pathways and loss of p53 tumor-suppressive pathways enables the cells to acquire an oncogenic phenotype.

**Supplementary figure 6**

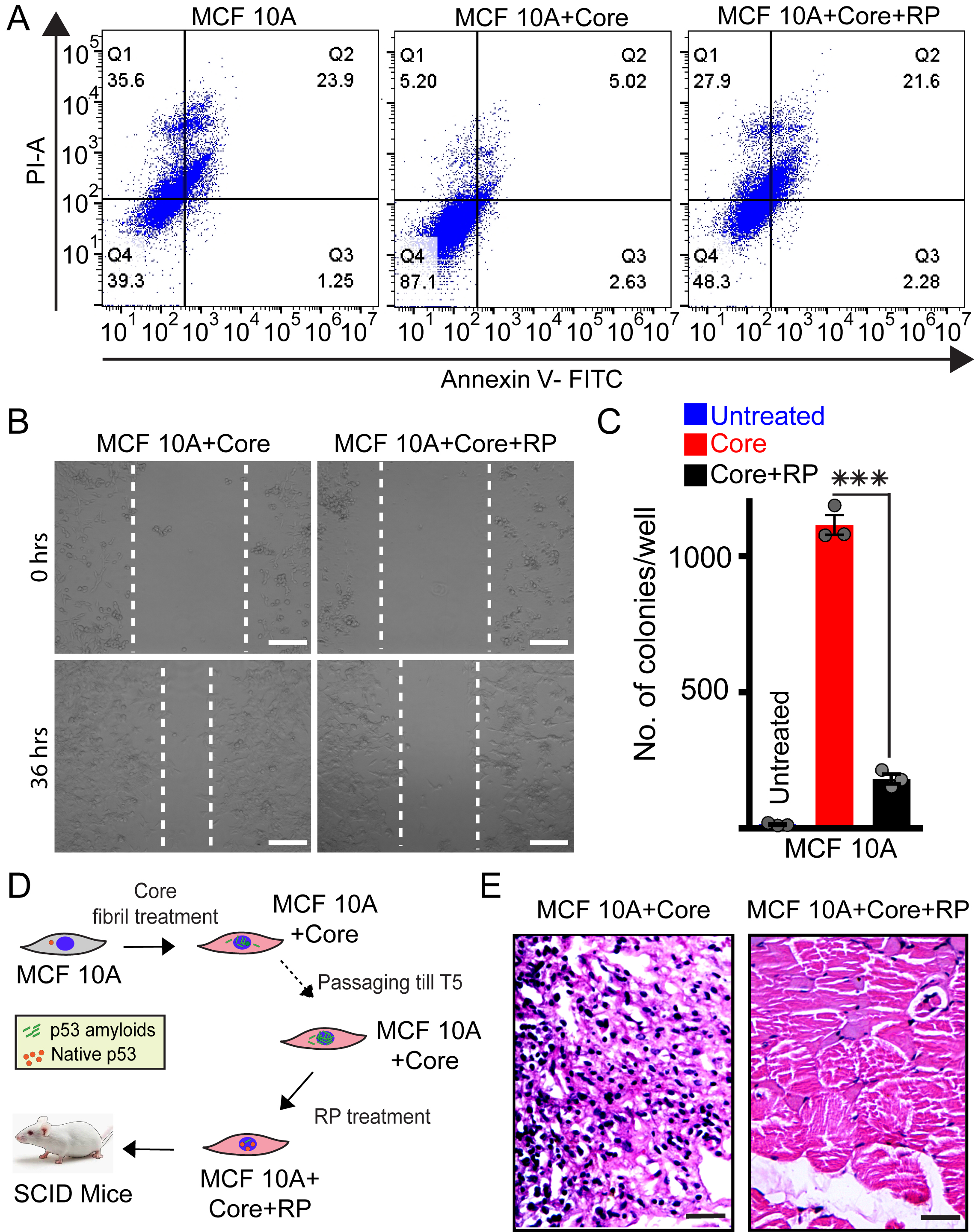
**Figure S6. Restoration of p53 activity upon disaggregation of p53 amyloids using rescue peptide (RP).** (A) Fibril transformed MCF 10A cells (at T5) were treated with rescue peptide (RP) at 10 µM concentration for 16 hrs to induce p53 disaggregation. Quantitative cell death assay using Annexin V and PI coupled with flow cytometric analysis showing RP treated cells with an increased apoptotic population (early and late combined) in comparison to only core fibril treated cells. n=3 independent experiments. (B) Cell motility using wound healing assay showing a decrease in the rate of cell migration by core fibrils treated cells (at T5) in presence of RP in comparison to only core fibril treated cells (at T5). Scale bars are 150 µm. (C) Quantification of colony number in the soft agar colony formation by core fibril transformed cells with or without RP treatment. The colonies formed per well for fibril treated and untreated cells were calculated and plotted. Fibril transformed cells (at T5) without rescue peptide treatment showed a large number of colonies, which drastically decreased upon RP treatment, indicating the rescue of p53 tumor-suppressive function. The values plotted represent mean ± s.e.m, n=3 independent experiments. Statistical significance is determined by one-way ANOVA followed by Bonferroni post hoc test; with p-value <0.0001. (D) Schematic representation of experimental setup for injection of rescue peptide treated cells. MCF 10A cells were treated with core fibrils and passaged till T5. Further, these cells were incubated in the presence of RP for 16 hrs and injected in SCID mice. MCF 10A core fibril treated cells at T5, without rescue peptide treatment, were injected as a control. These mice were monitored for tumor formation. (E) Hematoxylin and eosin (H & E) staining of tumor tissue formed by core fibril treated MCF 10A cells (at T5) injected in mice showing intense hematoxylin staining, suggesting the presence of hyperproliferative cells (left panel). RP treated cells (at T5) did not show tumor formation in mouse and the tissue H & E staining lacked the presence of hyperproliferative cells (right panel). Scale bars are 100 µm.

­**Supplementary figure 7**

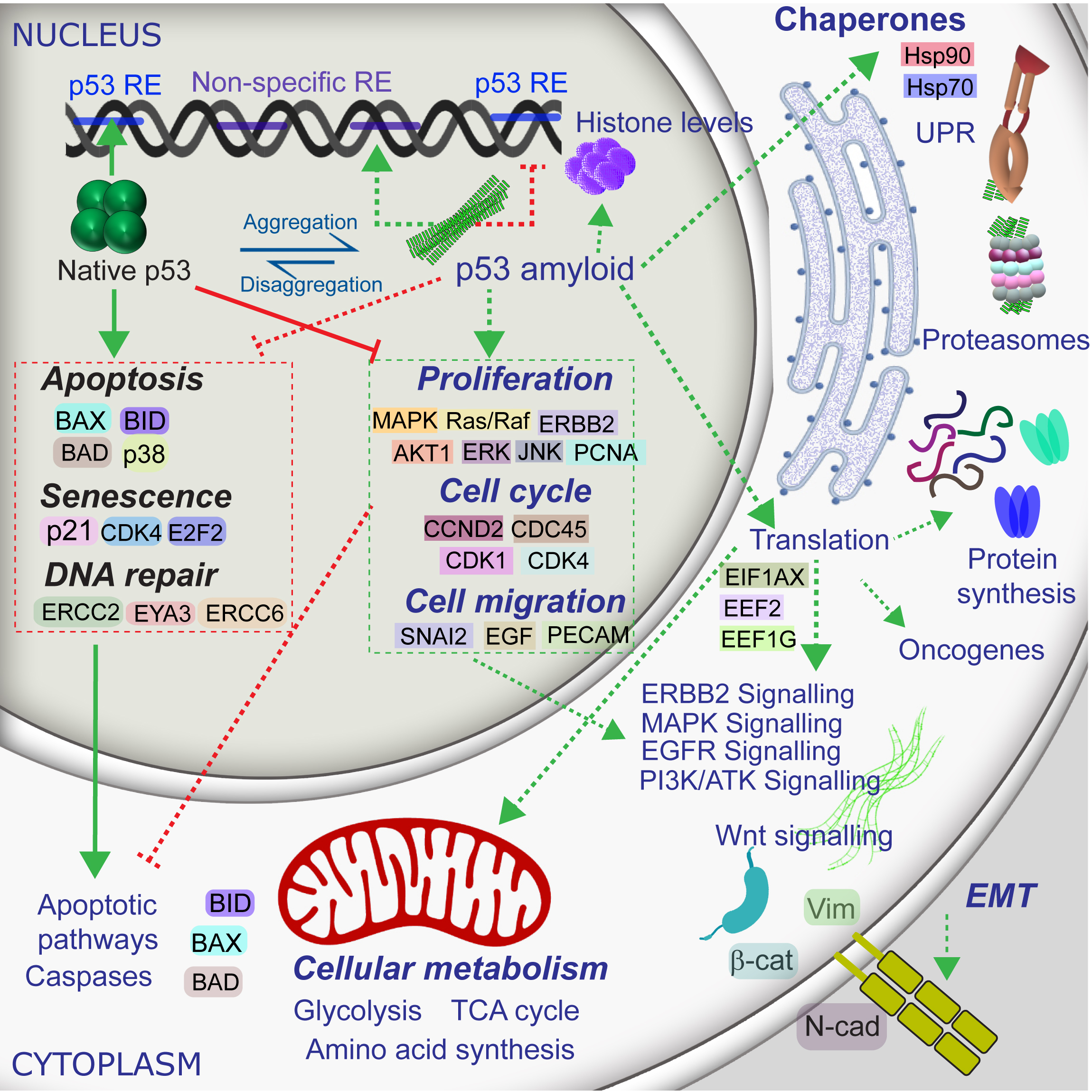

**Figure S7. Model of pathways affected by p53 amyloid formation in cells.** A mechanistic model of the cellular consequences due to p53 amyloid formation based on the microarray and proteomic analysis of cells harboring p53 aggregates. Solid and dotted arrows indicate processes affected by native p53 and amyloid form of p53, respectively. The red and green colour of arrows indicate negative regulation and positive regulation, respectively. p53 is a tumor suppressor protein, which regulates the expression of a plethora of downstream genes protecting the cells from the cancerous phenotype. Native p53 functions as a transcription factor in tetrameric form and binds to p53-specific response element (RE) to regulate the apoptotic, senescent and DNA repair pathways via multiple players. Native p53 also governs cell cycle and proliferation to maintain cellular homeostasis. Amyloid formation of p53 reverses its tumor-suppressive role and confers the gain of oncogenic function to the cells by modulation of cellular networks involved majorly in the cell cycle, DNA repair and cell proliferation. Pathways implicated in unfolded protein response, chaperones (Hsp 70, Hsp 90) and proteasomal machinery are highly upregulated due to amyloid formation. Due to p53 amyloid formation, genes and proteins involved in apoptotic and senescence pathways (such as Bax, Bid) are downregulated making the cell vulnerable to oncogenic transformation. Concurrently, as a downstream consequence, genes involved in cellular signaling, which induce proliferation and cell cycle (CDKs, MAPK, ErK, CDCs, Ras) are upregulated due to p53 amyloid formation. These pro-oncogenic genes confer growth, migratory and survival advantages to cells harboring p53 aggregates. Furthermore, the upregulation of proteins in translational and metabolic pathways supports the prolonged survival of these cells. p53 amyloid formation, thus, might affect the multistep process of malignant transformation, contributing differentially to cancer initiation, progression and metastasis.

**Supplementary table**

**Supplementary Table S1. List of primers used for real-time qPCR for gene expression study**

| **Gene** | **Forward Primer** | **Reverse Primer** |
| --- | --- | --- |
| GAPDH | 5’-CATTTTACGCTGATCCAGG-3’ | 3’-GGGTTCGAAATGAGGATG-5’ |
| p21 | 5’- AAGACCATGTGGACCTGT-3’ | 3’- GGTAGAAATCTGTCATGCTG-5’. |
| BAX | 5’-AACTGGACAGTAACATGGAG-3’ | 3’- TTGCTGGCAAAGTAGAAAAG-5’ |
| DDB2 | 5’-GAGCTTTGGAATCTCAGAATG-3’ | 3’-CAGGTCCCAAATTTTCACTG-5’ |
| MAPK1 | 5’-GAAGACTTATCTTGACCAGC-3’ | 3’-TCCATGGCACCTTATTTTTG-5’ |
| CCND2 | 5’-ACTTCATTGAGCACATCTTG-3’ | 3’-ACATGGCAAACTTAAAGTCG-5’ |

**Supplementary Table S1. List of primers used for real-time qPCR for gene expression study for various cell generations of fibril treated and untreated cells.** Primers used for analyzing the expression of genes linked with p53 pathways such as p21, pro-cancer genes (MAPK and CCND2) and anti-cancer genes (BAX, DDB2) were analyzed using real-time PCR (qRT-PCR) method (SYBR Green method).
